# Transmission of one predicts another: Apathogenic proxies for transmission dynamics of a fatal virus

**DOI:** 10.1101/2021.01.09.426055

**Authors:** Marie L.J. Gilbertson, Nicholas M. Fountain-Jones, Jennifer L. Malmberg, Roderick B. Gagne, Justin S. Lee, Simona Kraberger, Sarah Kechejian, Raegan Petch, Elliott Chiu, Dave Onorato, Mark W. Cunningham, Kevin R. Crooks, W. Chris Funk, Scott Carver, Sue VandeWoude, Kimberly VanderWaal, Meggan E. Craft

## Abstract

Identifying drivers of transmission prior to an epidemic—especially of an emerging pathogen—is a formidable challenge for proactive disease management efforts. We tested a novel approach in the Florida panther, hypothesizing that apathogenic feline immunodeficiency virus (FIV) transmission could predict transmission dynamics for pathogenic feline leukemia virus (FeLV). We derived a transmission network using FIV whole genome sequences, and used exponential random graph models to determine drivers structuring this network. We used these drivers to predict FeLV transmission pathways among panthers and compared predicted outbreak dynamics against empirical FeLV outbreak data. FIV transmission was primarily driven by panther age class and distances between panther home range centroids. Prospective FIV-based modeling predicted FeLV dynamics at least as well as simpler, often retrospective approaches, with evidence that FIV-based predictions captured the spatial structuring of the observed FeLV outbreak. Our finding that an apathogenic agent can predict transmission of an analogously transmitted pathogen is an innovative approach that warrants testing in other host-pathogen systems to determine generalizability. Use of such apathogenic agents holds promise for improving predictions of pathogen transmission in novel host populations, and could thereby provide new strategies for proactive pathogen management in human and animal systems.

## Introduction

Infectious disease outbreaks can have profound impacts on conservation, food security, and global health and economics. Mathematical models have proven a vital tool for understanding transmission dynamics of pathogens [1], but struggle to predict the dynamics of novel or emerging agents [2]. This is at least partially due to the challenges associated with characterizing contacts relevant to transmission processes. Common modeling approaches that assume all hosts interact and transmit infections to the same degree ignore key drivers of transmission. Such drivers can include specific transmission-relevant behaviors including grooming or fighting in animals [3], concurrent sexual partnerships in humans [4], or homophily [5], and result in flawed epidemic predictions [6,7]. Further, identifying drivers of transmission and consequent control strategies for any given pathogen is typically done reactively or retrospectively in an effort to stop or prevent further outbreaks or spatial spread (e.g. [8,9]). These constraints limit the ability to perform prospective disease management planning tailored to a given target population, increasing the risk of potentially catastrophic pathogen outbreaks, as observed in humans [10], domestic animals [11], and species of conservation concern (e.g., [12–14]).

A handful of studies have evaluated whether common infectious agents present in the healthy animal microbiome or virome can indicate contacts between individuals that may translate to interactions promoting pathogen transmission [15–22]. Such an approach circumvents some of the uncertainties associated with more traditional approaches to contact detection [6]. In these cases, genetic evidence from the transmissible agent itself is used to define between-individual interactions for which contact was sufficient for transmission to occur. Results of such studies show mixed success [15–18]. For example, members of the same household [19,20] or animals with close social interactions [21,22] have been found to share microbiota, but disentangling social mechanisms of this sharing is complicated by shared diets, environments, and behaviors [23].

These studies have, however, revealed ideal characteristics of non-disease inducing infectious agents (hereafter, *apathogenic agents*) for use as markers of transmission-relevant interactions. Such apathogenic agents should have rapid mutation rates to facilitate discernment of transmission relationships between individuals over time [24,25]. Furthermore, these agents should be relatively common and well-sampled in a target population, have a well-characterized mode of transmission that is similar to the pathogen of interest, and feature high strain alpha-diversity (local diversity) and high strain turnover [25,26]. RNA viruses align well with these characteristics [27] such that apathogenic RNA viruses could act as “proxies” of specific modes of transmission (i.e., direct transmission) and indicate which drivers underlie transmission processes. Such drivers, including but not limited to host demographics, relatedness, specific behaviors, or space use, would subsequently allow prediction of transmission dynamics of pathogenic agents with the same mode of transmission [25].

Here, we test the feasibility of this approach using a naturally occurring host-pathogen system to test if an apathogenic RNA virus can act as a proxy for direct transmission processes and subsequently predict transmission of a pathogenic RNA virus. Florida panthers (*Puma concolor coryi*) are an endangered subspecies of puma found only in southern Florida. We have documented that this population is infected by several feline retroviruses relevant to our study questions [28,29]. Feline immunodeficiency virus (FIVpco; hereafter, FIV) occurs in approximately 50% of the population and does not appear to cause significant clinical disease [28]. FIV is transmitted by close contact (i.e., fighting and biting), generally has a rapid mutation rate (intra-individual evolution rate of 0.00129 substitutions/site/year; [30]), and, as a chronic retroviral infection, can be persistently detected after the time of infection. Panthers are infected with feline leukemia virus (FeLV), also a retrovirus, which caused a well documented, high mortality outbreak among panthers in 2002-2004 [29]. FeLV infrequently spills over into panthers following exposure to infected domestic cats [31]. Once spillover occurs, FeLV is transmitted between panthers by close contact and results in one of three infection states: progressive, regressive, or abortive infection [29]. Progressive cases are infectious and result in mortality; regressive infections are unlikely to be infectious—though this is unclear in panthers— and recover [29,32,33]. Abortive cases clear infection and are not themselves infectious [32].

The objectives of this study were therefore: (1) to determine which drivers shape FIV transmission in Florida panthers, and (2) test if these drivers can predict transmission dynamics of analogously transmitted FeLV in panthers. Success of this approach in our model system would pave the way for testing similar apathogenic agents in other host-pathogen systems, thereby improving our understanding of drivers of individual-level heterogeneity in transmission, and consequently our ability to predict transmission dynamics of novel agents in human and animal populations.

## Methods

### Dataset assembly

We assembled an extensive dataset covering almost 40 years of Florida panther research and including panther sex and age class. A subset of the population is monitored using very high frequency (VHF) telemetry collars, with relocations determined via aircraft typically three times per week. Previous panther research has generated a microsatellite dataset for monitored panthers [34], and a dataset of 60 full FIV genomes (proviral DNA sequenced within a tiled amplicon framework in [35]). In addition, to augment observations from the 2002-04 FeLV outbreak [29], we leveraged an FeLV database which documents FeLV status (positive and negative) for 31 panthers from 2002-04 as determined by qPCR.

### FIV transmission inference

To determine drivers of FIV transmission, we first generated a “who transmitted to whom” transmission network using 60 panther FIV genomes collected from 1988 to 2011 (average of minimum annual panther counts across this period was 62.3 panthers; [36]). We used the program Phyloscanner [37] (see Figure 1 for workflow across all analyses), which assumes both within- and between-host evolution when inferring transmission relationships between sampled and even unsampled hosts [37]. Phyloscanner operates in a two step process, first inferring within- and between-host phylogenies in windows along the FIV genome. Then, using the within-host viral diversity gleaned from deep sequencing, Phyloscanner functionally performs ancestral state reconstruction to infer transmission relationships between hosts, outputting transmission trees or networks. For Phyloscanner’s step one, we used 150bp windows, allowing 25bp overlap between windows. To test sensitivity to this choice, we separately ran a full Phyloscanner analysis with 150bp windows, but without overlap between windows (supplementary methods). The tiled amplicon PCR approach used to generate our FIV genomic data biases for detection of one known variant, such that we did not expect detectable superinfections. In the second step of Phyloscanner, we therefore held the parameter which penalizes within-host diversity (k) equal to 0. We used a patristic distance threshold of 0.05 and allowed missing and more complex transmission relationships. Because we had uneven read depth across FIV genomes, we downsampled to a maximum of 200 reads per host. The output of the full Phyloscanner analysis was a single transmission network (hereafter, *main FIV network*), but see supplementary methods for details regarding analysis of the sensitivity of our results to variations in and summary across multiple transmission networks.

**Figure 1:**
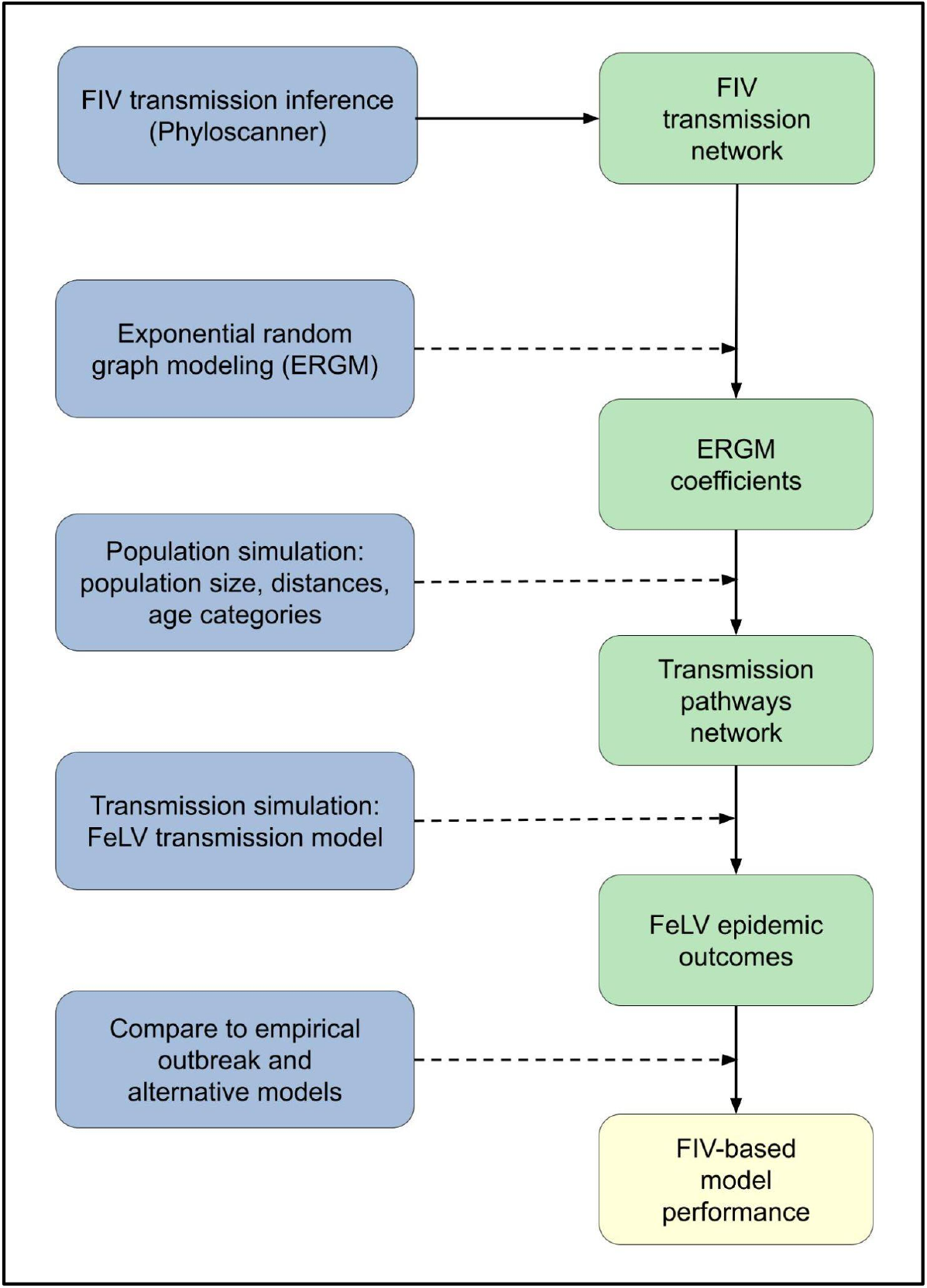
Conceptual workflow across all analysis steps. Processes are shown on the left in blue; specific outcomes are shown on the right in green; the final analysis outcome is in yellow at the bottom right. Solid lines show direct flows or outcomes. Dashed lines show processes acting on or in concert with prior outcomes: for example, exponential random graph modeling (ERGM) was performed using the FIV transmission network, and the combination of the two produced the ERGM coefficients outcome.

### Statistical analysis of FIV transmission networks

Phyloscanner transmission tree output suggests direction of transmission, but in our case, these results were often uncertain (see Results). To avoid putting undue emphasis on an uncertain direction of transmission, we simplified the transmission tree output to undirected, unweighted (binary) networks and performed statistical analysis of these networks using exponential random graph models (ERGMs; [38]). ERGMs model the edges in networks, with explanatory variables representing the potential structural drivers of the observed network [38]. By including network structural variables, ERGMs account for the inherent non-independence of network data. As such, we modeled “transmission relationships” (i.e., being connected in the transmission network) as a function of network structural variables and transmission variables we a priori expected to influence direct transmission processes in panthers. We considered several structural variables: an intercept-like edges term [38]; geometrically weighted edgewise shared partner distribution (*gwesp*; representation of network triangles); alternating k-stars (*altkstar*; representation of star structures); and 2-paths (2 step paths from *i* to *k* via *j*; [39]). In addition, we considered a suite of transmission variables (see supplementary methods for additional variable details): panther sex; age class (subadult or adult); pairwise genetic relatedness (panther microsatellite data from [34]); position of panther home range centroid (95% minimum convex polygon) or capture location (hereafter, *centroid*) relative to the major I-75 freeway (locations could be north or south of this east-west freeway); distance from centroid to nearest urban area (in km; USA Urban Areas layer, ArcGIS; [40]); pairwise geographic distance between centroids (log-transformed; Figure S1); and pairwise home range overlap (utilization distribution overlap indices of 95% bivariate normal home range kernels; [41,42]).

Because ERGMs are prone to degeneracy with increasing complexity [38], we first performed forward selection for network structural variables, followed by forward selection of dyad-independent variables, while controlling for network structure. Model selection was based on AIC and goodness of fit, and MCMC diagnostics were assessed for the final model (supplementary methods). ERGMs were fit with the *ergm* package [43] in R (v3.6.3; [44]).

### Panther population simulations

We next simulated FeLV transmission through a network representing panthers during the 2002-04 FeLV outbreak where network edges represented likely transmission pathways based on ERGM-identified predictors of FIV transmission (*FIV-based model*). Hereafter, a *full-simulation* includes both simulation of the panther population with its likely transmission pathways and simulation of FeLV transmission within that population. Below, we describe the process for a single simulation, but these procedures were repeated for each full simulation.

We first based the simulated population size on the range of empirical estimates from 2002-2004 (Table S1; [36]). Additional characteristics of the simulated population included those identified as significant variables in the ERGM analysis: age category and pairwise geographic distances between panther home range centroids (see Results). We randomly assigned age categories to the simulated population based on the proportion of adults versus subadults. Age proportions were based on age distributions in the western United States [45], which qualitatively align with the historically elevated mean age of the Florida panther population [46]. Pairwise geographic distances for the simulated population were generated by randomly assigning simulated home range centroids based on the distribution of observed centroids on the landscape (supplementary methods).

We then used ERGM coefficients to generate network edges among the simulated panther population using the *ergm* package in R [43]. The FIV transmission network spanned 15 years of observations and represents a subset of the actual contact network, as it includes only those interactions that resulted in successful transmission. We therefore had a high degree of uncertainty regarding the appropriate network density for our simulations. To manage this uncertainty, we constrained density (ratio of existing edges to all possible edges) in our network simulations across a range of parameter space (net_dens, Table S1).

### Simulation of FeLV transmission on FIV-based networks

The next step in each full simulation was to model FeLV transmission through the network generated from FIV predictors of transmission. FeLV transmission was based on a stochastic chain binomial process on the simulated network, following a modified SIR compartmental model (Figure 2). Simulations were initiated with one randomly selected infectious individual and proceeded in weekly time steps. Transmission simulations lasted until no infectious individuals remained or until 2.5 years, whichever came first.

**Figure 2:**
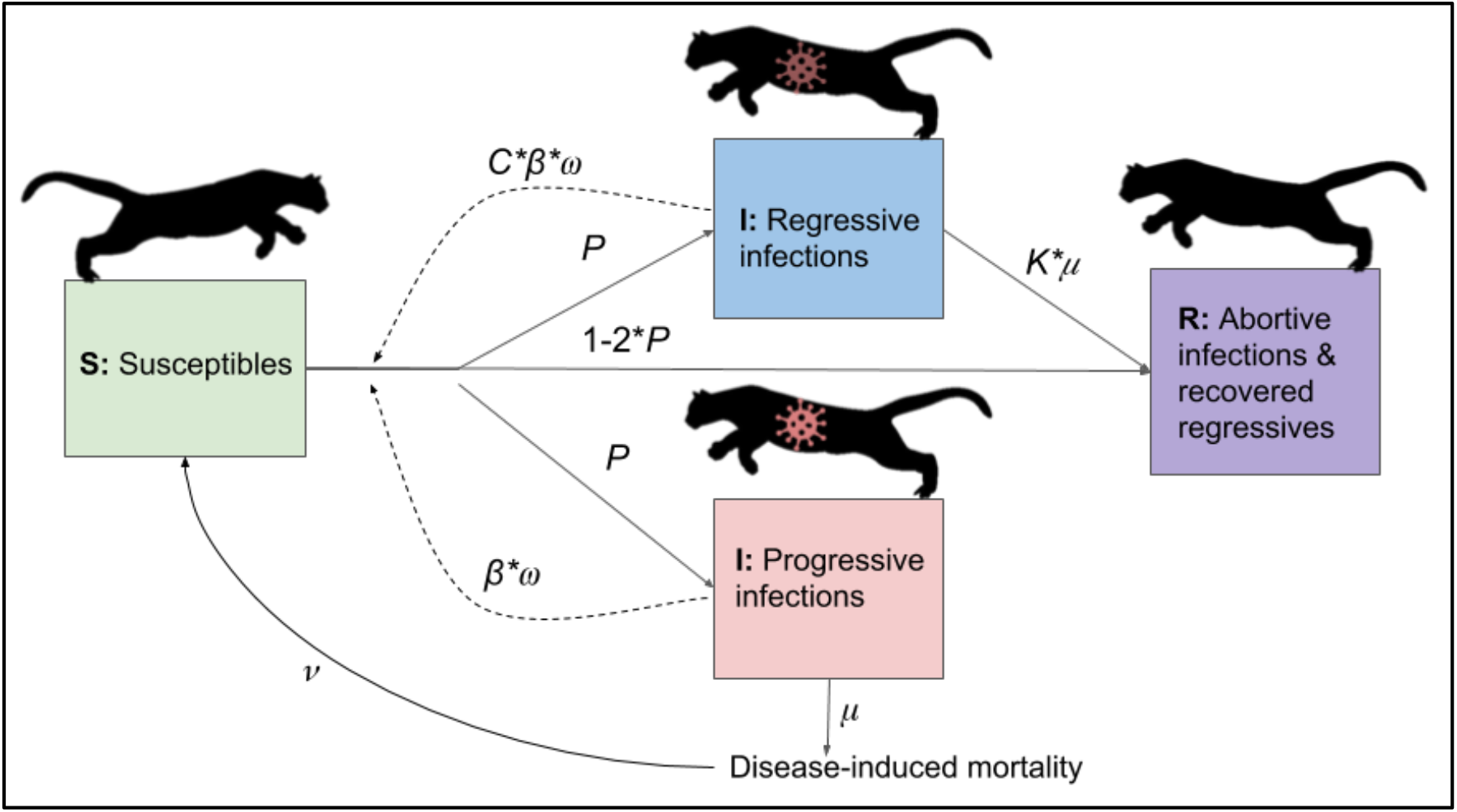
Diagram of flows of individuals between compartments in transmission model. Virus icons indicate infectious states, with the regressive infection icon darkened to represent reduced or uncertain infectiousness of this class. Note: a vaccination process was also included in the transmission model, but is not shown for simplicity. With vaccination, susceptibles could be vaccinated, and vaccinated individuals subsequently infected as with susceptibles, but with an additional probability of (1-ve). See Table S1 for definitions of parameters.

Transmission was dependent on the following (Figure 2; see Table S1 for parameter definitions): (1) existence of an edge between two individuals, (2) the dyad in question involving a susceptible and infectious individual, and (3) a random binomial draw based on the probability of transmission given contact (*β*). In addition, *Puma concolor* generally have low expected weekly contact rates [47]; we therefore included a weekly contact probability, represented as a random binomial draw for contact in a given week (*ω*).

Upon successful transmission, infectious individuals were randomly assigned to one of three outcomes of FeLV infection [29]. *Progressive* infections (probability *P*) are infectious (*β*), develop clinical disease, and die due to infection (*μ*). *Regressive* infections (also probability *P*) recover from infection (*K*\**μ*, where *K* is a constant) and, having entered a state of viral latency, are not considered at risk of FeLV reinfection [29,48]. Using model assumptions derived from known patterns of FeLV infection in domestic cats, regressive individuals are not infectious [29], but given ongoing uncertainty, we included some transmission from regressives (*C*β*, where *C* is a constant). *Abortive* infections (probability 1-2*P*) are never infectious, clearing infection and joining the recovered class. While the duration of immunity in abortive cases has not been studied in panthers, because abortive cases clear infection through a strong immune response and develop anti-FeLV antibodies, reinfection with FeLV is considered extremely unlikely [48].

A vaccination process was included in simulations as panthers were vaccinated against FeLV during the historical FeLV outbreak starting in 2003. Vaccination occurred at a rate, *τ*, and applied to the whole population, as wildlife managers are unlikely to know if a panther is susceptible at the time of capture or darting. However, only susceptible individuals transitioned to the vaccinated class (i.e. vaccination failed in non-susceptibles). Because panthers were vaccinated in the empirical outbreak with a domestic cat vaccine with unknown efficacy in panthers, we allowed vaccinated individuals to become infected in transmission simulations by including a binomial probability for vaccine failure (1-vaccine efficacy, *ve*, Table S1).

The panther population size remained roughly static through the course of the FeLV outbreak [36]. We therefore elected not to include background mortality, but did include infection-induced mortality. To maintain a consistent population size, we therefore included a birth/recruitment process. Because FIV-based simulated networks drew edges based on population characteristics, we treated births as a “respawning” process, in which territories vacated due to mortality were reoccupied by a new susceptible at rate, *ν*. This approach allowed us to maintain the ERGM-based network structure and is biologically reasonable, as vacated panther territories are unlikely to remain unoccupied for long.

### Comparison of simulation predictions to observed FeLV outbreak

To evaluate the performance of our FIV-based model, we also predicted FeLV transmission dynamics using three alternative models: random networks, home range overlap-based networks, and a well-mixed model. The random networks model used Erdős-Rényi random networks, matching network densities from the FIV-based model (Table S1), but otherwise allowing edges to occur between any pairs of individuals. Overlap-based networks were generated using the degree distributions of panther home range overlap networks from 2002-2004 and simulated annealing with the R package *statnet* ([49]; supplementary methods). For both random and overlap-based networks, FeLV transmission was simulated as in the FIV-based simulations. The well-mixed model was a stochastic, continuous time compartmental model (Gillespie algorithm), with rate functions aligning with the chain binomial FeLV transmission probabilities (see supplementary methods).

We performed transmission simulations for all *model types* (FIV-based, overlap-based, random, and well-mixed) across a range of reasonable parameter space (Table S1), using a Latin hypercube design (LHS) to generate 150 parameter sets that efficiently sampled parameter space [50,51]. For each parameter set and model type, we performed 50 simulations (30,000 total). In each simulation, we recorded the number of mortalities and the duration of outbreaks, which were each summarized (medians) across each parameter set. For each model type, we determined if each parameter set’s predicted median mortalities, duration of outbreaks, and abortive cases were within a reasonable range based on the observed FeLV outbreak (5-20 mortalities, 78-117 week duration, at least 5 abortive infections; [29]). If so, a parameter set was deemed “feasible” for that model type. Ranges were used to account for uncertainty in observations and population size in this cryptic species (supplementary methods). To test for differences in the frequency of feasible FeLV predictions between model types, we fit a binomial generalized linear mixed model (GLMM), assuming a logistic regression with “feasible” (vs “unfeasible”) as the outcome, model type as a predictor variable, and a random intercept for parameter set.

We tested for spatial clustering of cases in the observed FeLV outbreak by leveraging our database of qPCR-based FeLV status. We performed a local spatial clustering analysis of FeLV cases and controls using SaTScan (50% maximum, circular window; [52]). A SaTScan analysis seeks to identify clusters of cases in which the observed cases within a particular cluster exceed random expectation; this analysis reports the observed/expected ratio and radius of any significant clusters. In addition, we performed a global cluster analysis with Cuzick and Edward’s test (global cluster detection with case-control data) in the R package *smacpod* (1, 3, 5, 7, 9, and 11 nearest neighbors; 999 iterations; [53–55]). To determine if simulated FeLV cases likewise demonstrated spatial clustering, we repeated SaTScan local cluster analysis and Cuzick and Edward’s tests (at 3, 5, and 7 nearest neighbors) with feasible FIV-based simulation results. To determine if detected clustering in FIV-based simulations was simply based on our respawning protocol, we also performed both spatial analyses with feasible overlap-based simulation results as a “negative control.” Because the overlap-based model was not spatially explicit, we assigned the same geographic locations to nodes in the overlap-based networks from the corresponding FIV-based networks.

To determine if certain transmission parameters were important for feasible outcomes, we performed post hoc random forest variable importance analyses for each of the four model types with “feasible” as the binary response variable (using the R package randomForest [56,57]; see supplementary results).

## Results

### FIV transmission network analysis

In the main FIV network, Phyloscanner inferred 42 potential transmission relationships (edges) between 19 individuals (nodes; network density = 0.25), after removing 9 edges that were between individuals known not to be alive at the same time (Figure 3). Panther FIV genomes missing from the transmission network were those for whom transmission relationships could not be inferred by Phyloscanner (see Discussion). ERGM results for the main FIV network identified triangle (*gwesp*) and star structures (*altkstar*) as key structural variables, and age category and log transformed pairwise geographic distance as key transmission variables (Tables 1, S2). Though altkstar was not statistically significant, inclusion of this variable contributed to improved AIC and goodness of fit outcomes. Adults were more likely to be involved in transmission events (but see discussion of sample size limitations) and inferred transmission events were more likely between individuals which were geographically closer to each other. The fitted model showed reasonable goodness of fit (Figures S2-3).

**Figure 3:**
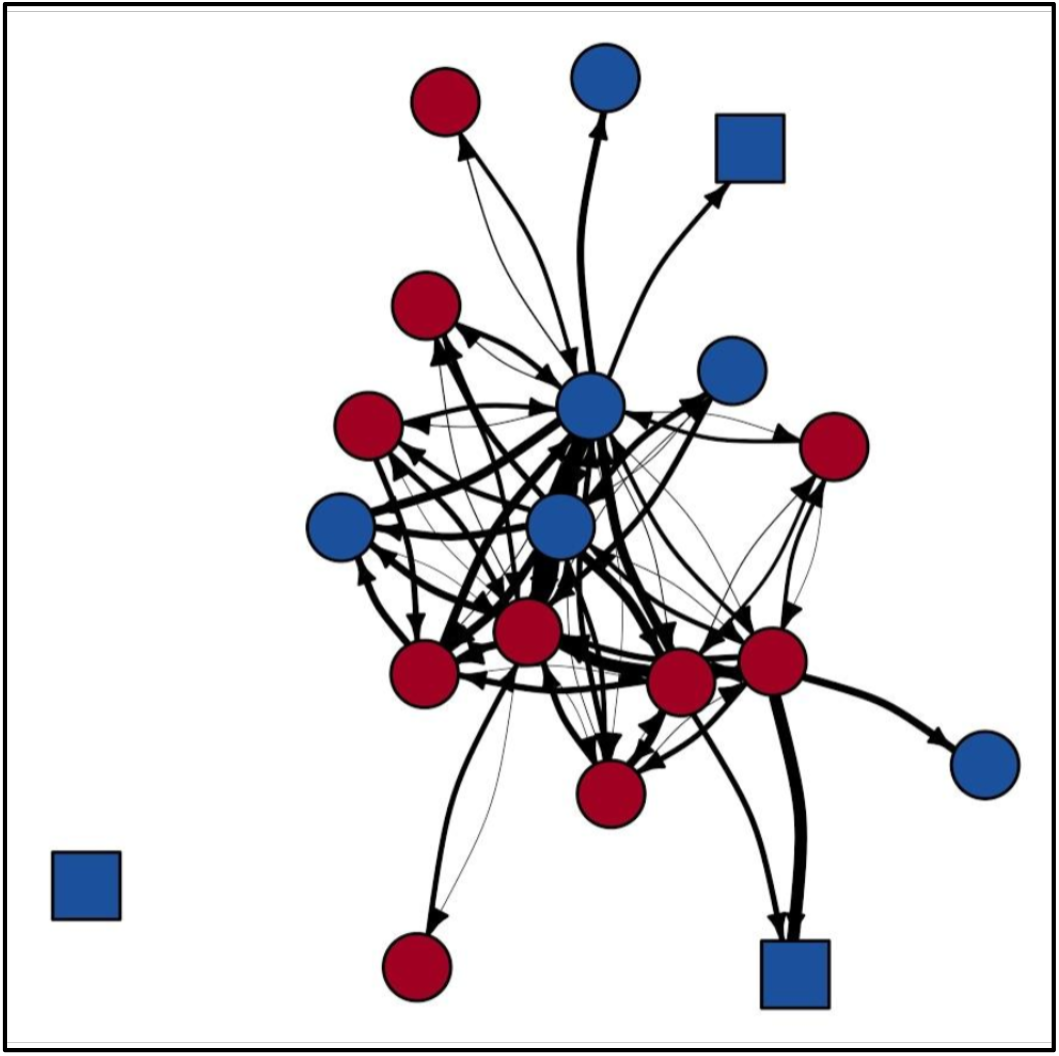
Phyloscanner-derived main FIV transmission network. Node shape indicates panther age class (square = subadult; circle = adult). Node color indicates panther sex (blue = male; red = female). Edge weight represents Phyloscanner tree support for each edge (thicker edge = increased support); for visualization purposes, edges are displayed as the inverse of the absolute value of the log of these support values. While pictured as a directed and weighted network, statistical analyses used binary, undirected networks.

**Table 1:**
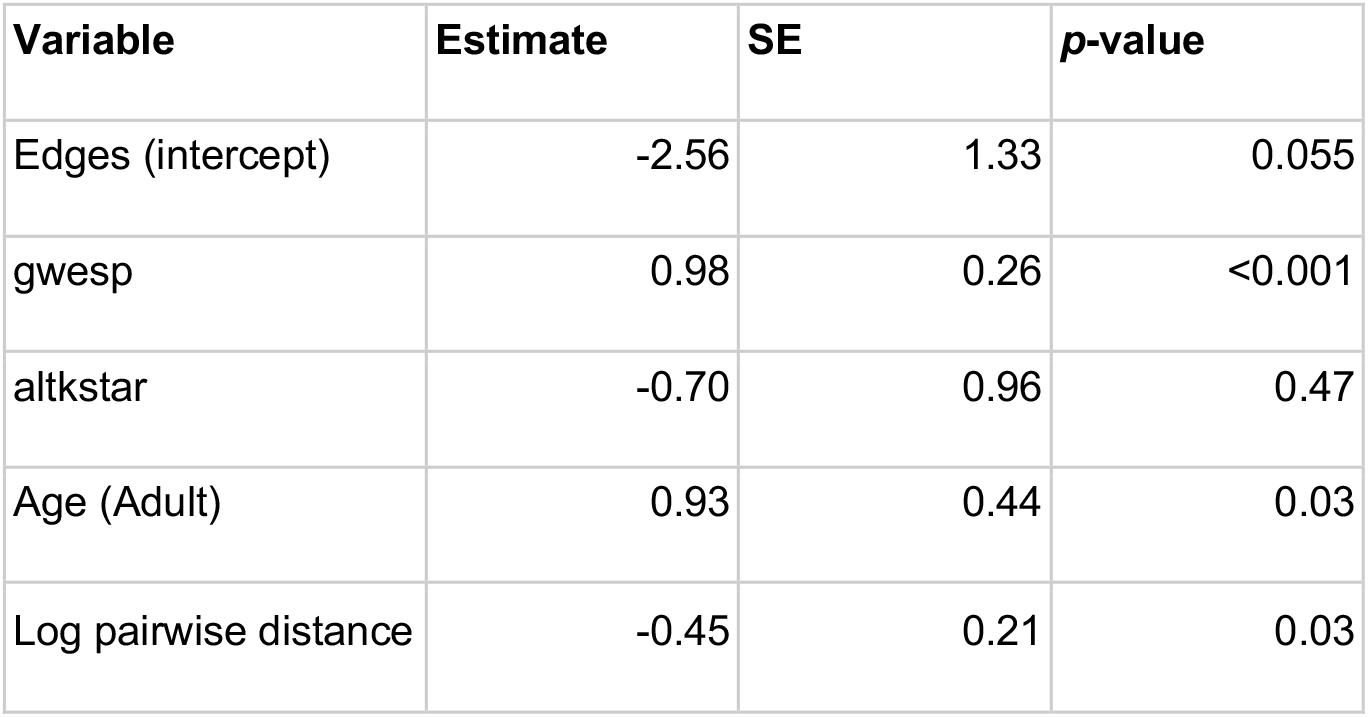
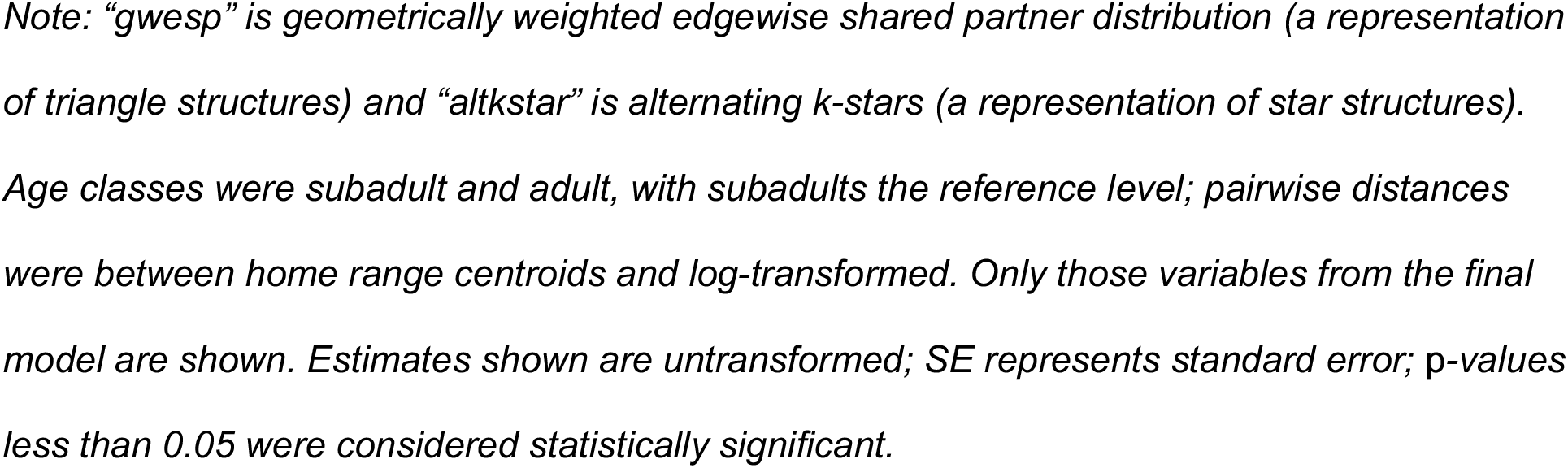
Main FIV transmission network exponential random graph model results.

### FeLV simulations

About 9% of parameter sets across all model types were classified as feasible (Figures S6-S7). The FIV-based model had the highest odds of feasibility, though this difference did not achieve statistical significance (Table 2). SaTScan analysis of observed FeLV status found weak evidence of local spatial clustering (two clusters detected, but not statistically significant with *p*=0.165 and 0.997, respectively; Figure S5). Cuzick and Edward’s tests found evidence of global clustering at 3, 5, and 7 nearest neighbor levels (test statistic T_k_ where *k* is number of nearest neighbors considered: T_3_ = 20, *p* = 0.049; T_5_ = 32, *p* = 0.028; T_7_ = 43, *p* = 0.023). Feasible parameter sets from both the FIV-based and overlap-based models produced some evidence of local and global spatial clustering of simulated FeLV cases (Figures 4, S8). However, the FIV-based model better captured the size and strength of predicted local clusters (SaTScan radius and observed/expected cases, respectively; Figure 4) and was moderately better at capturing global spatial patterns (Figure S8).

**Table 2:**
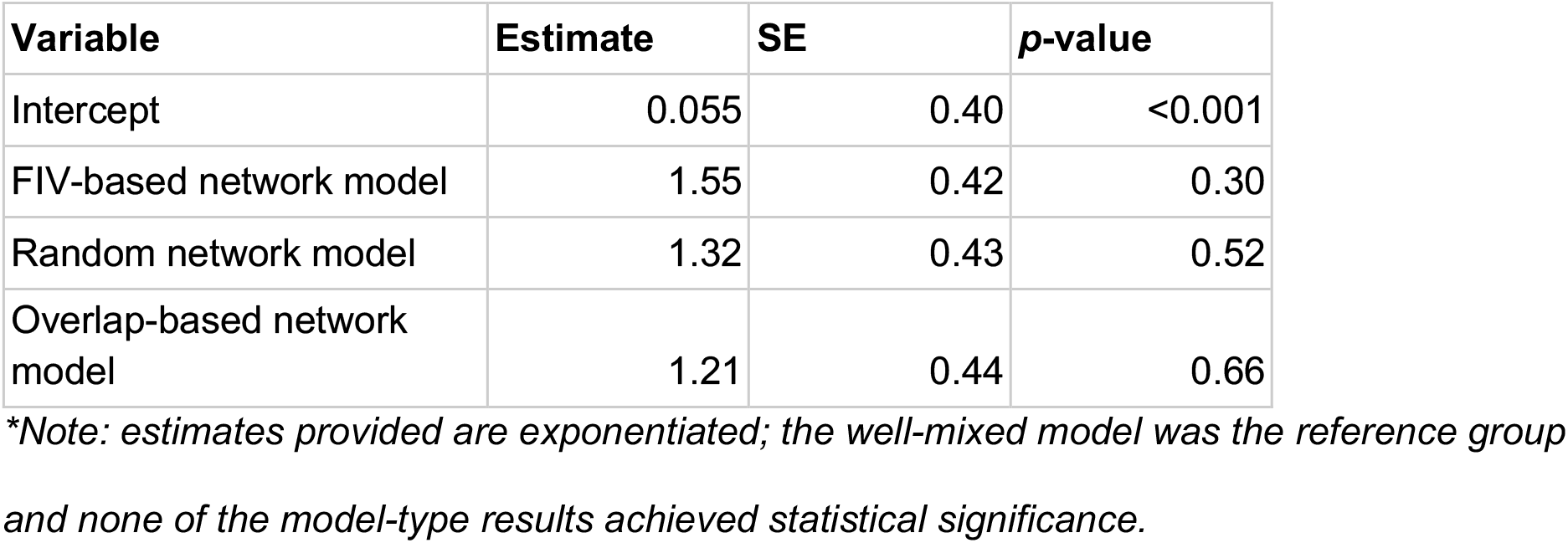
Fixed effects results from model-type performance GLMM*.

| Variable | Estimate | SE | p-value |
| --- | --- | --- | --- |
| Intercept | 0.055 | 0.40 | <0.001 |
| FIV-based network model | 1.55 | 0.42 | 0.30 |
| Random network model | 1.32 | 0.43 | 0.52 |
| Overlap-based network model | 1.21 | 0.44 | 0.66 |

*\*Note: estimates provided are exponentiated; the well-mixed model was the reference group and none of the model-type results achieved statistical significance.*

**Figure 4:**
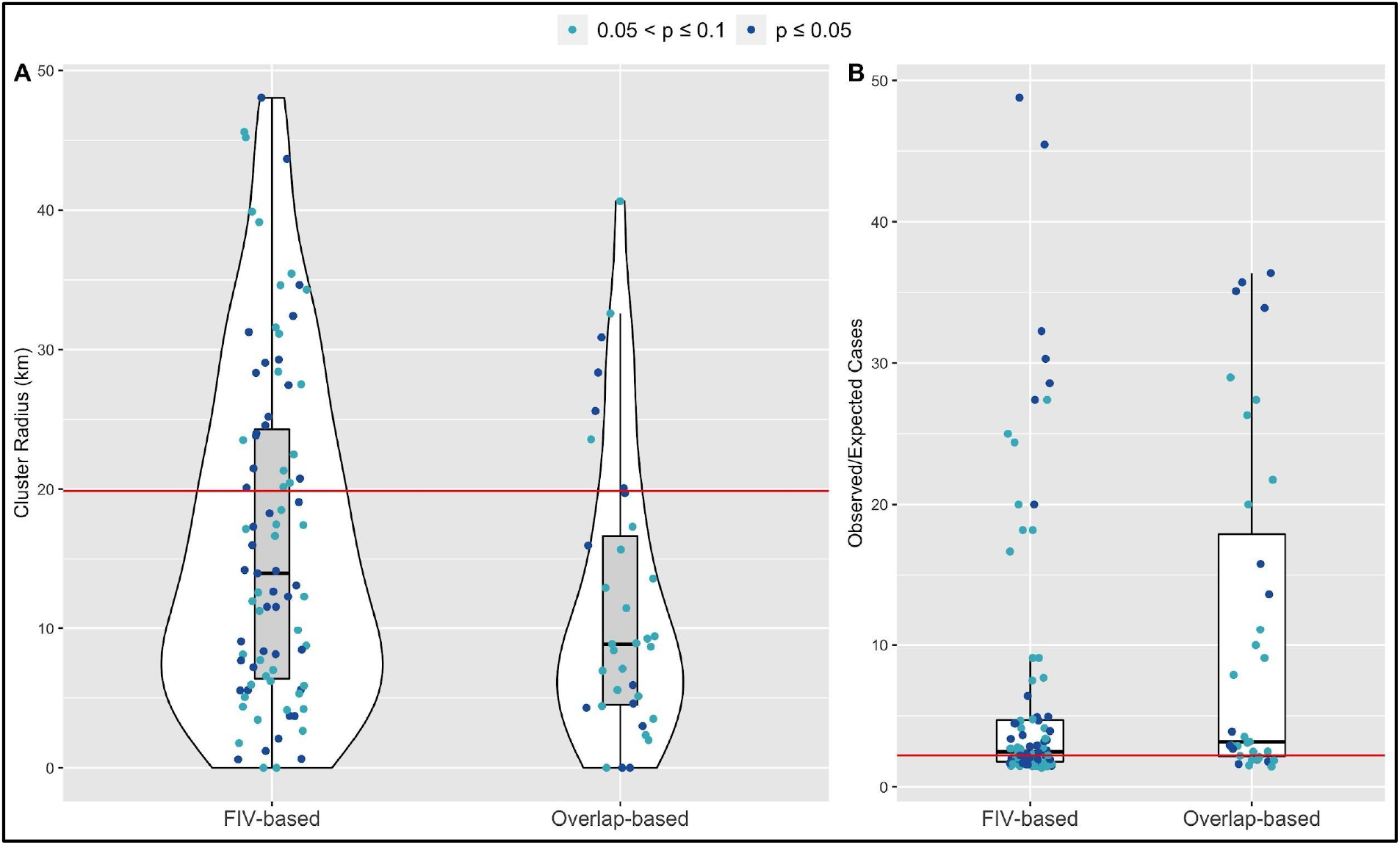
SaTScan cluster analysis for feasible FIV-based and overlap-based network simulations show stronger agreement for the FIV-based model, compared to the overlap-based model, between empirical observations (red horizontal lines) and model predictions for (A) FeLV cluster size and (B) Observed/Expected FeLV cases associated with the top detected cluster. Shown are feasible simulation results in which at least one cluster was detected with p-values less than or equal to 0.1; further, only the results from the top cluster are shown.

The post-hoc random forest analyses typically showed poor balanced accuracy and area under the curve (AUC) results. However, the parameter shaping transmission from regressively infected individuals (C), consistently showed support for weak to moderate transmission from regressives (i.e., C = 0.1 or 0.5; Figure S11).

## Discussion

In this study we develop a new approach whereby we leverage knowledge of transmission dynamics of a common apathogenic agent to prospectively predict dynamics of an uncommon and virulent pathogen. Our approach was distinctly different from simpler models we tested, as the apathogenic (FIV)-based approach focused on underlying drivers or mechanisms of transmission and could be used to prospectively identify predictors of transmission and develop disease control plans prior to an outbreak of a virulent pathogen (FeLV). We found that FIV transmission in panthers is primarily driven by distance between home range centroids and age class, and that our prospective FIV-based approach predicted FeLV transmission dynamics at least as well as simpler or more reactive approaches. While we do not propose that this apathogenic agent approach could accurately predict exactly when, where, and to whom transmission might occur, our results support the role of apathogenic agents as novel tools for prospectively determining sources of individual-level heterogeneity in transmission and consequently improving proactive disease management.

### An FIV-based model captures FeLV transmission dynamics

We found that our network model based on drivers of FIV transmission produced FeLV outbreak predictions consistent with the observed FeLV outbreak. The FIV-based approach performed at least as well as simpler models, per our GLMM analysis, with evidence that FIV better predicted the observed spatial dynamics for FeLV transmission. A key difference between the FIV-based approach and other spatially explicit methods is that FIV allowed us to determine the importance of spatial dynamics prospectively and then translate to predictions of FeLV transmission, rather than relying on retrospective FeLV spatial analyses. Furthermore, while more complex potential drivers of transmission (e.g., host relatedness or assortative mixing by age or sex) were not found to be important for FIV transmission, these may yet be key for structuring transmission in other systems. Simpler model types like random networks or metapopulation models may struggle to make transmission predictions that incorporate these drivers of transmission-relevant contact. The predictive power we observed here using an apathogenic virus could thus open new opportunities to determine behavioral and ecological drivers of individual-level heterogeneity in the context of pathogen transmission, and even shape proactive epidemic management strategies for pathogens such as FeLV.

Our network statistical analysis (ERGMs) determined that pairwise geographic distances and age category structure FIV transmission in the Florida panther. These findings are well supported by panther and FIV biology, providing confidence in the functioning of our apathogenic agent workflow. For example, panthers are wide-ranging animals but maintain home ranges, and this appears to translate to increased transmission between individuals that are close geographically. This finding is supported by the tendency for FIV phylogenies to show distinct broad [58] and fine scale [59] geographic clustering in *Puma concolor*. Further, specifically among Florida panthers, spatial autocorrelation of FIV exposure status was previously found to approach statistical significance [60]. The wide-ranging nature of puma appears to limit geographic clustering of many infectious agents [60], with FIV a notable exception to this pattern. In addition, because FIV is a persistent infection, we would expect cumulative risk of transmission to increase over an individual’s lifetime and adults would consequently be involved in more transmission events. The low number of subadult individuals in our dataset, however, means that this finding must be interpreted with some caution.

With these ERGM results in mind, key components of the success of our apathogenic agent approach are likely that (1) FIV is a largely species-specific virus with transmission pathways closely matching intraspecific transmission of FeLV, and (2) both FIV and FeLV, perhaps unusually for infectious agents of puma, display spatial clustering of infection. Here, FIV fundamentally acted as a proxy for close, direct contact in panthers, and could consequently determine drivers of such contacts. If, for example, FIV also exhibited strong vertical or environmental transmission, we would no longer expect the predictive success for FeLV we observed here. This consideration highlights the importance of careful apathogenic agent selection when attempting to predict pathogen transmission, a workflow which must now be tested in additional host-pathogen systems. For example, the mixed results when using commensal agents to identify close social relationships in other systems [15–18,21,22] highlights that some host-apathogenic agent combinations will work better than others for determining drivers of transmission. Within our study, Phyloscanner struggled to elucidate transmission relationships between many of our FIV genomes, likely due to unusually low genetic diversity among our FIV isolates, or our use of proviral DNA (which has lower diversity than circulating RNA) [35]. While the drivers of transmission we identified are biologically reasonable, we may have lacked the power to identify more complex relationships (e.g., homophily) due to the low number of individuals in our transmission network.

Elaborating on agent selection suggestions outlined in the introduction, we propose that apathogenic agent selection should carefully consider agent genetic diversity within a target population—not just expected diversity based on typical mutation rates, as in our case—and favor those agents with high diversity to facilitate transmission inference. We also propose that apathogenic agents should represent the timescales of transmission for the pathogen of interest. For example, FeLV produces long, slow epidemics, such that FIV transmission relationships may be most representative across the longer timescales we evaluated here. In contrast, short, acute pathogen epidemics would likely best be represented by apathogenic agent transmission over shorter timescales. Our results highlight that, perhaps most importantly, an apathogenic agent should have a well characterized mode of transmission that closely matches transmission of the pathogen of interest, as this was likely key to our success with FIV and FeLV. Future research should determine how divergent an apathogenic agent may be from a pathogen of interest while still predicting transmission dynamics.

While few parameter sets in our simulations were classified as feasible, this appears to be predominantly the result of the wide range of parameter space explored through our LHS sampling design. This limitation was fundamentally due to uncertainties in FeLV transmission parameters, and is representative of the uncertainties experienced in predicting transmission of emerging or understudied pathogens. Infectiousness of regressives, the number of introductions of FeLV to the panther population, and the duration of FeLV infection in panthers were all important sources of uncertainty in our models. All three of these dynamics can have significant impacts on the duration of a simulated epidemic, allowing an epidemic to continue to “stutter” along at low levels [61], much as was observed in the empirical FeLV outbreak. Our *post hoc* random forest analysis provided some evidence of weak transmission from regressive individuals, but this finding would need to be validated with additional research, as it is in stark contrast to FeLV dynamics in domestic cats. Reducing uncertainties in these three key dynamics would significantly narrow the range of our predictions, and even assist in ongoing management efforts for FeLV in endangered panthers. The effect of transmission parameter uncertainties underscores the importance of linking laboratory and model-based research to generate more accurate transmission forecasts [62].

### Limitations and future directions

The suite of tools for inferring transmission networks from infectious agent genomes is rapidly expanding [24]. In this study, we used the program Phyloscanner as it maximized the information from our deep sequencing viral data. However, our FIV sequences were generated within a tiled amplicon framework [35,63], which biases intrahost diversity and limits viral haplotypes [64]. Phyloscanner was originally designed to analyze RNA from virions and not proviral DNA, as we have done here. We have attempted to mitigate the effects of these limitations by analyzing several different Phyloscanner outputs to confirm consistency in our results, and by using only binary networks to avoid putting undue emphasis on transmission network edge probabilities, as these are likely uncertain. Further, our primary conclusions from the transmission networks—that age and pairwise distance are important for transmission—are biologically plausible and supported by other literature, as discussed above. Nevertheless, future work should evaluate additional or alternative transmission network inference platforms.

In addition, ERGMs assume the presence of the “full network” and it is as yet unclear how missing data may affect transmission inferences [38]. ERGMs are also prone to degeneracy with increased complexity and do not easily capture uncertainty in transmission events, as most weighted network ERGM (or generalized ERGM) approaches have been tailored for count data (e.g., [65]). ERGMs may therefore not be the ideal solution for identifying drivers of transmission networks in all systems. Alternatives may include advancing dyad-based modeling strategies [66], which may more easily manage weighted networks and instances of missing data.

Our FIV-based approach required extensive field sampling, and many disciplines from viral genomics through simulation modeling. However, with increasing availability of virome data and even field-based sequencing technology, our approach may become more accessible with time. Further, the predictive benefits seen here, while needing further testing and validation, could become a key strategy for proactive pathogen management in species of conservation concern, populations of high economic value (e.g. production animals), populations with infrequent pathogen outbreaks that make targeted surveillance more difficult, or populations at high risk of spillover, all of which may most benefit from rapid, efficient epidemic responses.

### Conclusions

Here, we integrated genomic and network approaches to identify drivers of FIV transmission in the Florida panther. This apathogenic agent acted as a marker of close, direct contact transmission, and was subsequently successful in predicting the observed transmission dynamics of the related pathogen, FeLV. Further testing of apathogenic agents as markers of transmission and their ability to predict transmission of related pathogens is needed, but holds promise for revolutionizing proactive epidemic management across host-pathogen systems.

## Acknowledgements

Thanks to M. Michalska-Smith, K. Worsley-Tonks, J. Mistrick, and S. N. Hart for key feedback. This research was supported by the National Science Foundation (DEB-1413925, 1654609, and 2030509). MLJG was supported by the Office of the Director, National Institutes of Health (NIH T32OD010993), the University of Minnesota Informatics Institute MnDRIVE program, and the Van Sloun Foundation. JLM was supported by the ACVP/STP Coalition for Veterinary Pathology Fellows and the Linda Munson Fellowship for Wildlife Pathology Research. The content is solely the responsibility of the authors and does not necessarily represent the official views of the National Institutes of Health. Puma icon by Freepik at Flaticon.com.

## Data Availability

Data and R code for replication of simulations and analyses is available on GitHub (https://github.com/mjones029/FIV_FeLV_Transmission) and upon acceptance will be archived at Zenodo.

