## Supplementary Materials for "Transmission of one predicts another: Apathogenic proxies for transmission dynamics of a fatal virus"

---

---

### Methods

#### *Phyloscanner*

For feline immunodeficiency virus (FIV) transmission inference, we used Phyloscanner to generate a main transmission network. For step one of Phyloscanner analysis, we used 150bp windows, allowing 25bp overlap between windows. For step two, we held k, which penalizes within-host diversity, equal to 0. We used a patristic distance threshold of 0.05 and allowed missing and more complex transmission relationships. Because we had uneven read depth across FIV genomes, we downsampled to a maximum of 200 reads per host. To test sensitivity of our results to variations in Phyloscanner output, we also generated two *summary FIV networks*, varying the degree of window overlap in the first step of Phyloscanner analysis. With Phyloscanner step one set to 25bp overlap, we generated four additional FIV transmission networks, but kept only those edges that were found in at least two of these four networks. We repeated this process with Phyloscanner step one set to 0bp overlap, again keeping only those edges found in at least two of four resulting transmission networks. We then removed any edges between individuals that were known not to be alive at the same time, based on panther monitoring data. This process resulted in our main FIV network, and the two additional summary FIV networks.

#### *Statistical analysis of FIV transmission networks*

To determine factors structuring FIV transmission networks, we used an exponential random graph modeling approach (ERGM). As described in the main text, ERGMs model the edges in networks, with explanatory variables representing the potential structural drivers of the observed network [1]. We considered a suite of network structural variables (also called dyad-dependent variables; see main text), which can account for the non-independence in network data, and our key transmission predictors of interest (dyad-independent variables). When evaluating unweighted (binary), undirected networks, dyad-independent variables operate like explanatory variables in a logistic regression, and can include both node-level variables (e.g., node age or sex) and edge-level variables (e.g., genetic distance). Further, node-level variables can be evaluated as continuous or categorical variables (e.g., are males involved in more transmission events?), but can also test for difference or matching relationships which capture homophily within the network (e.g. do more transmission events occur between male/female dyads or same-sex dyads?). In keeping with ERGM analysis terminology, categorical node variables are referred to as *node factor*, continuous as *node covariate*, matching relationships as *node matching*, mixing relationships (not constrained to homophily) as *node mixing*, and continuous edge variables as *edge covariate* [1,2].

Among the dyad-independent variables we examined in our ERGM analysis (see main text), we evaluated panther sex and age class as both node factors and node mixing variables. For panther age class, subadults were classified as individuals between the ages of 6 months and two years, and adults classified as individuals over two years of age.

Our ERGM analysis also included several spatial variables. Home range centroids were used in the generation of several of these variables and were determined by first estimating 95% minimum convex polygon (MCP) home ranges for telemetry-monitored panthers. To estimate these MCP home ranges, we used only those telemetry data collected in the 12 months after an individual’s initial capture, and only for those individuals with at least 30 relocations in that time period. MCPs were generated with the *adehabitatHR* package in R [3], and centroids were calculated using the *rgeos* package [4]. Our priority was to capture the range of panther-occupied landscape, so we also incorporated point locations for individuals without at least 30 telemetry relocations. For these point locations, we prioritized using the location of an individual at capture; if this information was not available, we instead used the telemetry relocation collected closest to the date of capture. For the main FIV network, this approach resulted in 11 point locations from MCP centroids, 7 were capture locations, and 1 was a relocation closest to capture date. Hereafter, these locations are referred to as *centroids*.

Major roadways have been shown to alter puma (*Puma concolor*) movement in North America [5], so we hypothesized that panthers would be more likely to transmit to panthers on the same side of Florida's major I-75 freeway as themselves. Our ERGM analysis therefore included a node-matching variable for location of panthers' centroids north versus south of the I-75 freeway (which runs east-west through panther habitat), defined as latitude 26.15. We further hypothesized that panthers closer to urban areas would face greater competition for resources and therefore be involved in more transmission events due to increased fighting behaviors. We therefore also examined a node covariate term for distance to the nearest urban area (in km). We used the "near table" function in ArcGIS to determine the distance of each centroid to the closest urban area edge, defining urban areas using the USA Urban Areas layer publicly available in ArcGIS (Census 2010 Urbanized Areas and Clusters; [6]). We included pairwise geographic distances between panthers using distances between centroids (in km), and log-transformed this edge covariate for ERGM analysis (Figure S1). Lastly, we hypothesized that panthers with overlapping home ranges would be more likely to transmit to each other, so we included a spatial overlap edge covariate based on the pairwise utilization distribution overlap indices (UDOI) of 95% bivariate normal home range kernels [7], using the default settings in the *adehabitatHR* package in R [3]. Of note: UDOI assesses overlap between utilization distributions, such that the home range kernels used in this analysis are distinct from the MCP home ranges used to estimate centroid locations.

For our pairwise relatedness variable (see main text), we used previously published microsatellite data [8]. One individual in the FIV transmission networks lacked microsatellite data, but had known pedigree sibling relationships with other individuals in the transmission networks [9]. In order to preserve available data (ERGMs cannot operate with missing data), we interpolated sibling relatedness values for this individual using mean relatedness values from other known sibling pairs. Non-sibling relationships for this individual were conservatively interpolated at population mean relatedness, functionally assuming no relatedness.

Goodness of fit for ERGMs was performed using the *ergm* package in R [10]. We evaluated fit for degree distribution, geodesic distance, and triad census (“degree”, “distance,” and “triadcensus” terms, respectively). Model selection included evaluation of AIC and improvement to goodness of fit, using these terms.

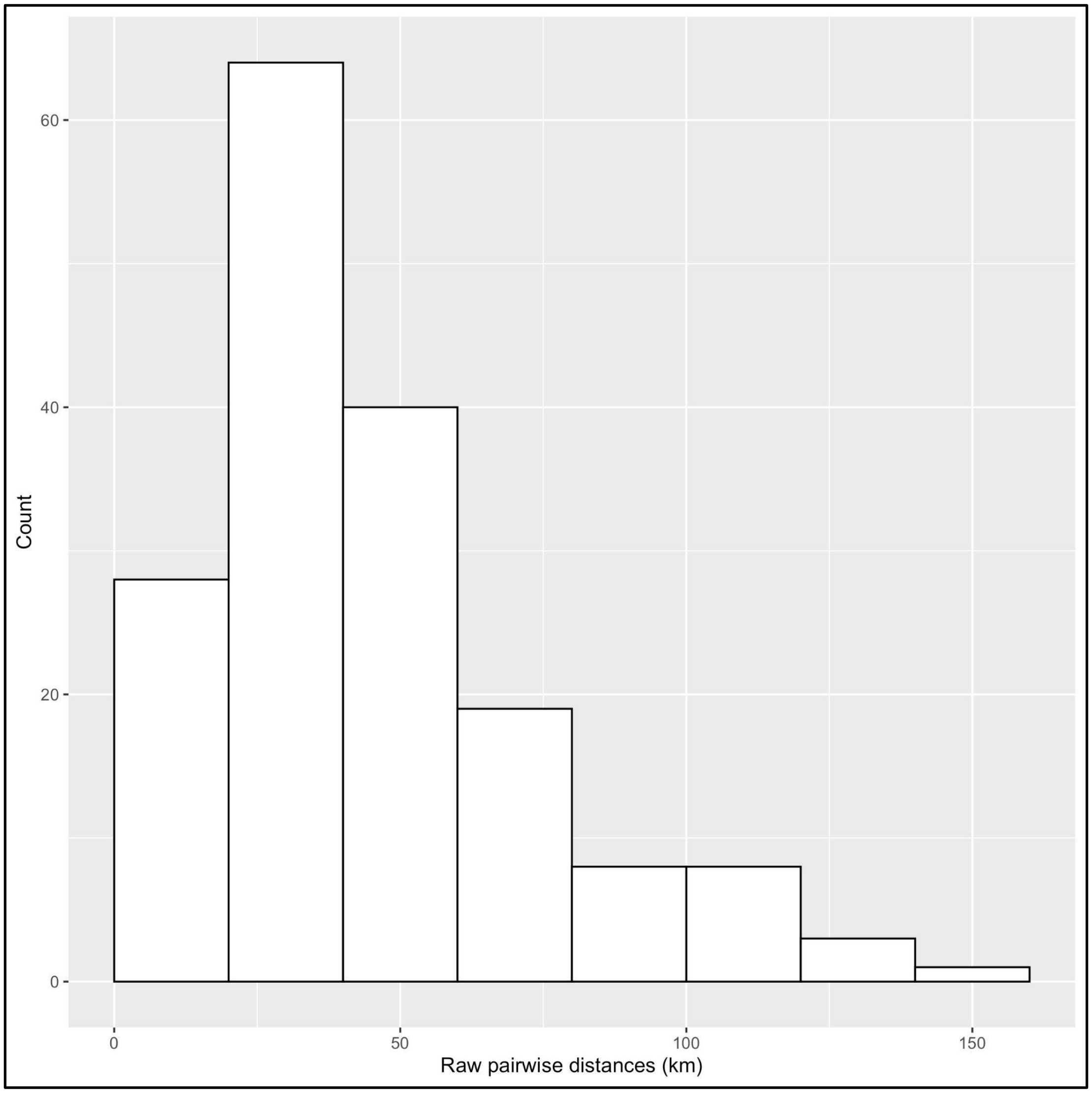

**Figure S1:** Histogram of raw (untransformed) pairwise distances between centroids.

*Panther centroid simulation*

Because pairwise geographic distances were found to be significant in ERGM analysis of FIV transmission networks (see main text), in order to simulate potential transmission pathways among panthers, we also had to simulate these geographic pairwise distances. We did so by simulating home range centroids based on the empirical panther population. Simulated centroids were generated by plotting the observed MCP centroids from 2002-2004; the polygon encompassing these centroids was then split into 70 quadrats, and simulated centroids were randomly drawn from these quadrats, according the proportion of the observed population that was found within each quadrat. This functionally kept much of the heterogeneity in distribution of home range centroids across panther habitat. Pairwise distances were then calculated between simulated centroids and log transformed, as was done to calculate pairwise distances for the original ERGM analysis (see main text).

*Overlap-based networks*

To compare FeLV transmission predictions from FIV-based networks against simpler model types, we generated spatial overlap-based networks, through which we also simulated FeLV transmission (see main text). To do so, we first generated networks of utilization distribution overlap index (UDOI) spatial overlap (with 95% bivariate normal kernel) among collared panthers in each year from 2002-2004 (three total networks; [3]), considering an edge to exist if UDOI was greater than 1. We calculated the degree distribution from each of these networks, and fit a negative binomial distribution to the degree distribution for each year using the *fitdistrplus* package in R [11]. We took the means of the parameter values for the resulting three negative binomial distributions to create a single "summary" negative binomial distribution. We simulated new overlap-based networks using this summary negative binomial to draw degree distributions, and then used simulated annealing [12,13] to generate random networks based on the drawn distributions. Because simulated overlap-based networks were informed by degree distributions, they were not spatially explicit, but represent data typically available in long-term wildlife monitoring studies.

**Table S1: Network and transmission simulation parameters**

| Parameter | Definition | Range | Reference |
|---|---|---|---|
| Pop_size | Population size | 80-120 | [14] |
| Adult_prop | Proportion adults versus subadults | 0.82-0.99 | [15] |

| | | | |
|---|---|---|---|
| Net_dens | Simulated network density | 0.05-0.15 | NA |
| $\beta$ | Probability of transmission from progressives, given effective contact | 0.17-0.29 | [16] |
| *C* | Constant multiplier for probability of transmission from regressives, given effective contact | 0, 0.1, 0.5, 1 | NA |
| $\omega$ | Weekly probability of contact | 0.1-0.4 | [17] |
| $\mu$ | Weekly probability of death from progressive infection | 1/18, 1/26* | [18] |
| *K* | Constant multiplier for weekly probability of recovery from regressive infection | 0.5, 1 | NA |
| $\nu$ | Weekly probability of territory repopulation ("respawn rate") | 1/12-1/4 | NA |
| $\tau$ | Weekly probability of vaccination | 0.5-1 | NA |
| *ve* | Probability of vaccine efficacy | 0.4-1 | [18] |
| *P* | Proportion randomly assigned to each of the progressive and, regressive states† | 0.25 | [18] |

*Note: Parameter gives parameter symbols or abbreviations; definition gives the description for each parameter. Range shows the continuous range or discrete values sampled from in simulations, with references giving literature supporting ranges or values. *We tested a lower death rate (prolonged duration of infection) due to the low number of observed panther cases and the generally longer infection duration in domestic cats [19]. †The proportion randomly assigned to the abortive state was therefore 1-2P (0.50).*

*Gillespie algorithm*

We also compared FeLV transmission predictions from FIV-based networks to predictions from a homogeneous mixing model: in this case, a Gillespie algorithm (stochastic, time-to-event model). This model was specified in order to align with the chain binomial network model specifications (Figure 2; main text), resulting in the following rate functions:

Susceptibles infection rate = $\omega*\beta*\text{Net\_dens}*S(I_p + C*I_r)$

Vaccinates infection rate = $\omega*\beta*(1-ve)*\text{Net\_dens}*V(I_p + C*I_r)$

Progressives mortality rate = $\mu I_p$

Recovery rate = $\mu*K*I_r$

Respawn rate = $\nu D$

Vaccination rate (after one simulation year) = $\tau*S/N$

In the above rate functions, N is the total population, S is susceptibles, $I_p$ is progressively infected individuals, $I_r$ is regressively infected individuals, V is vaccinated individuals, and D marks unoccupied territories after death of the prior occupant and prior to "respawning" (as in network models). All other parameters are as in Table S1. Of special note, Net_dens represents the density of networks from network transmission models, and here functions as a population size-scaled contact rate. This contact rate is further modified by the weekly probability of contact, $\omega$, as was done in network models (see main text). The vaccination rate is scaled by population size as vaccination was applied to the whole population in both network and Gillespie models, but only susceptibles could transition from susceptible to vaccinated.

*FeLV spatial analyses*

To test for spatial clustering of FeLV in the observed panther outbreak, we used a dataset of FeLV qPCR results (n = 31), in which 12 individuals tested positive. We used a circular window with a maximum spatial cluster of 50% of the population at risk. In addition, we used the same data to test for global clustering using Cuzick and Edward's test with the *smacpod* package in R [20,21]. Here, we evaluated nearest neighbor levels (k) of 1, 3, 5, 7, 9 and 11, and used 999 iterations for inference. We used these same parameterizations for

SaTScan and Cuzick and Edward’s analyses of FeLV simulations, with the exception that we only evaluated k = 3, 5, and 7 for simulated data (see main text).

*FeLV prediction target ranges*

When comparing FeLV simulation predictions against observations from the historical outbreak, we used several “target” ranges for outbreak duration and the number of progressive infections. More specifically, the empirical outbreak is considered to have occurred from July 1, 2002 - June 30, 2004 (104 weeks), but due to uncertainty in the precise duration of the historical outbreak, we considered a simulated duration of 78-117 weeks to be “on target.” During the observed outbreak, 5 individuals were documented with progressive (or transient) infection. Panthers are cryptic, difficult-to-observe animals, resulting in uncertainty in detection of all progressive infections and full population size at the time. We therefore considered 5-20 progressive infections in simulations to be on target.

While our primary focus was progressive infections, we also included an expectation that at least 5 individuals were abortive infections. Empirically, these individuals were the most numerous, but as they were not clinically ill, abortive infections were less likely to be detected in normal panther management; we therefore did not include an upper bound for this target.

### Results

*FIV transmission network inference*

Phyloscanner FIV network results indicated some dissimilarities between single runs. As reported in the main text, the main FIV network included 19 nodes with 42 edges (network density = 0.25) after removing 9 edges that were between individuals known not to be alive at the same time (Figure 3 of main text). The summary transmission network allowing scanning window overlap included 20 nodes with 43 edges (network density = 0.23), and the summary network without window overlap included 20 nodes with 35 edges (network density = 0.18; after 8 and 6 edges removed, respectively, due to dates known alive).

*Supplementary ERGM results*

Results from the best ERGM for each FIV network are given in Table S2. Of note, the best ERGM for the main FIV network showed reasonable goodness of fit (Figures S2-3). ERGM results for the two summary networks were comparable to the main FIV network (Table S2). The key difference was that the summary network with no window overlap did not find log transformed pairwise geographic distances to be a significant variable, though this fitted model showed evidence of degeneracy. To further confirm consistency of our ERGM findings, we performed a *post hoc* analysis with simulated random networks (see below; Figure S4).

**Table S2: Best ERGM results for each FIV transmission network**

| Transmission network | Variable | Estimate | SE | *p*-value |
|---|---|---|---|---|
| Main FIV network | Edges (intercept) | -2.56 | 1.33 | 0.055 |
| | gwesp | 0.98 | 0.26 | <0.001 |
| | altkstar | -0.70 | 0.96 | 0.47 |
| | Age (Adult) | 0.93 | 0.44 | 0.03 |
| | Log pairwise distance | -0.45 | 0.21 | 0.03 |
| Summary network with window overlap | Edges (intercept) | -0.15 | 1.48 | 0.92 |
| | gwesp | 1.03 | 0.31 | <0.001 |
| | altkstar | -3.51 | 1.22 | 0.004 |
| | Age (Adult) | 1.36 | 0.61 | 0.02 |
| | Log pairwise distance | -0.63 | 0.22 | 0.004 |
| Summary network without window overlap | Edges (intercept) | -2.76 | 1.33 | 0.038 |
| | gwesp | 1.03 | 0.32 | 0.001 |

| | | | | |
|---|---|---|---|---|
| | altkstar | -2.17 | 0.99 | 0.029 |
| | Age (Adult) | 1.03 | 0.57 | 0.073 |

*Note: “gwesp” is geometrically weighted edgewise shared partner distribution (a representation of triangle structures) and “altkstar” is alternating k-stars (a representation of star structures). Age classes were subadult and adult, with subadults the reference level; pairwise distances were between home range centroids and log-transformed. Only those variables from the final model are shown. Estimates shown are untransformed; SE represents standard error;* p-values *less than 0.05 were considered statistically significant.*

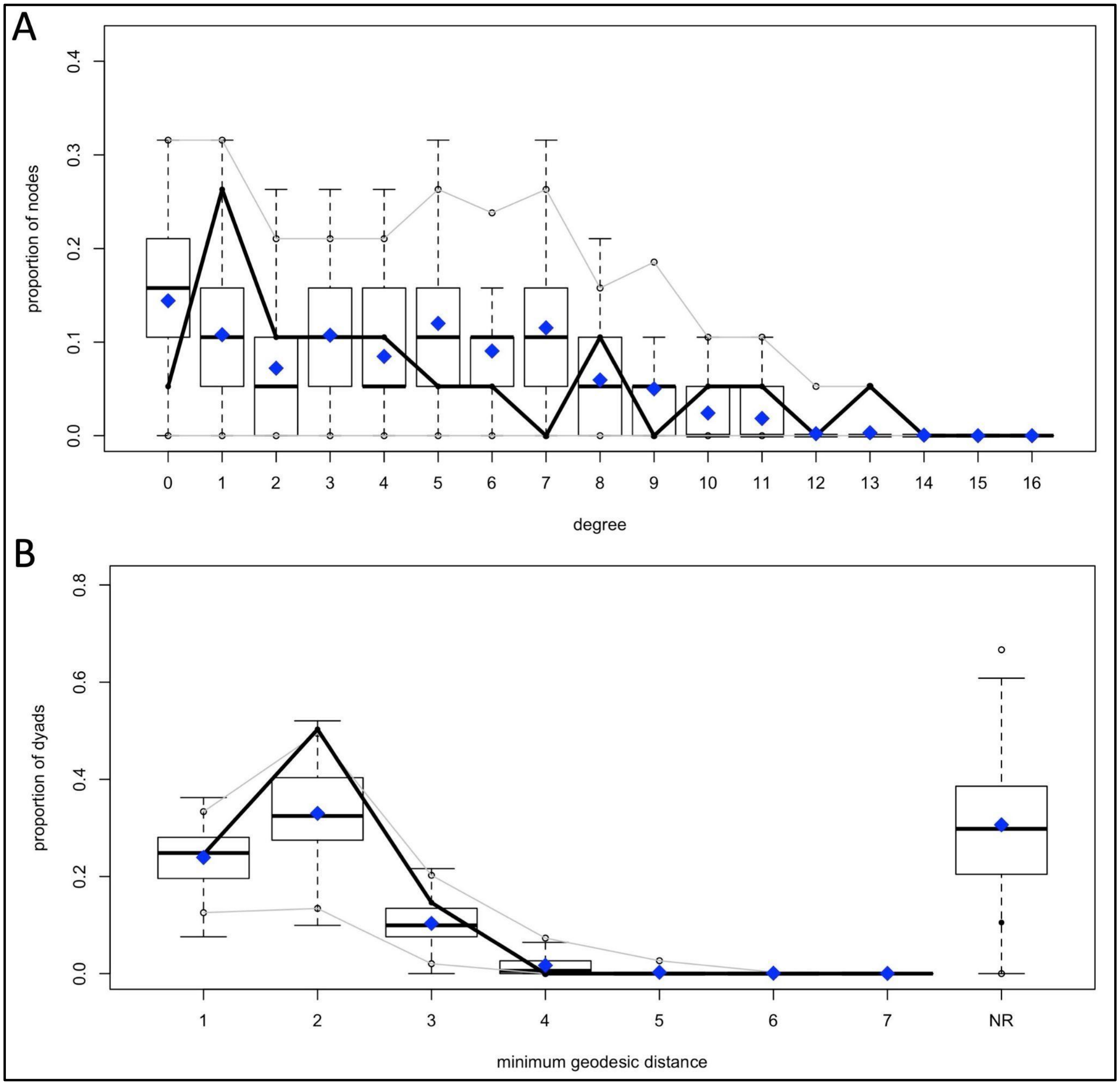

***Figure S2:*** *With Figure S3, best ERGM for main FIV network showed reasonable goodness of fit across standard goodness of fit metrics: (A) degree, (B) minimum geodesic distance. Boxplots show model predictions; solid black lines observations.*

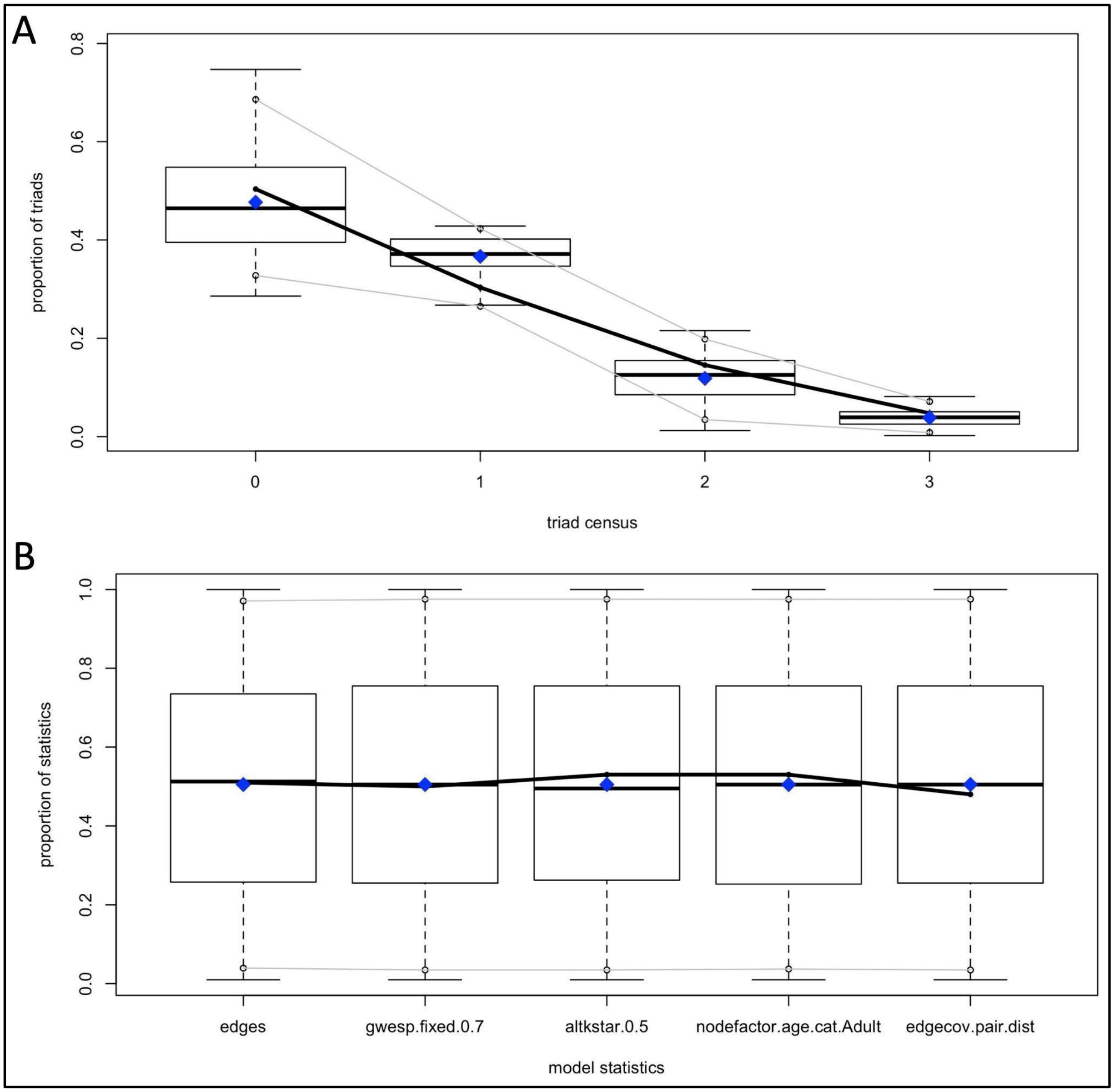

***Figure S3:*** *With Figure S2, best ERGM for main FIV network showed reasonable goodness of fit across standard goodness of fit metrics: (A) triad census, (B) model statistics. Boxplots show model predictions; solid black lines observations.*

*Post hoc random network ERGM analysis*

Because there were some differences between ERGM results from the three FIV transmission networks (main text; Table S2), we performed a *post hoc* random network analysis to determine the consistency of our results against "null" random networks. Using the same panthers and descriptive data from the main FIV transmission tree, we rewired this transmission network as an Erdős-Rényi random network of the same density and fit an ERGM with the same variables from our main ERGM. We repeated this procedure 50 times, recording variable coefficients with each iteration. We then compared the distribution coefficients from simulated random networks to those from our three ERGMs, finding strong consistency among our ERGM coefficients relative to those from random networks (Figure S4).

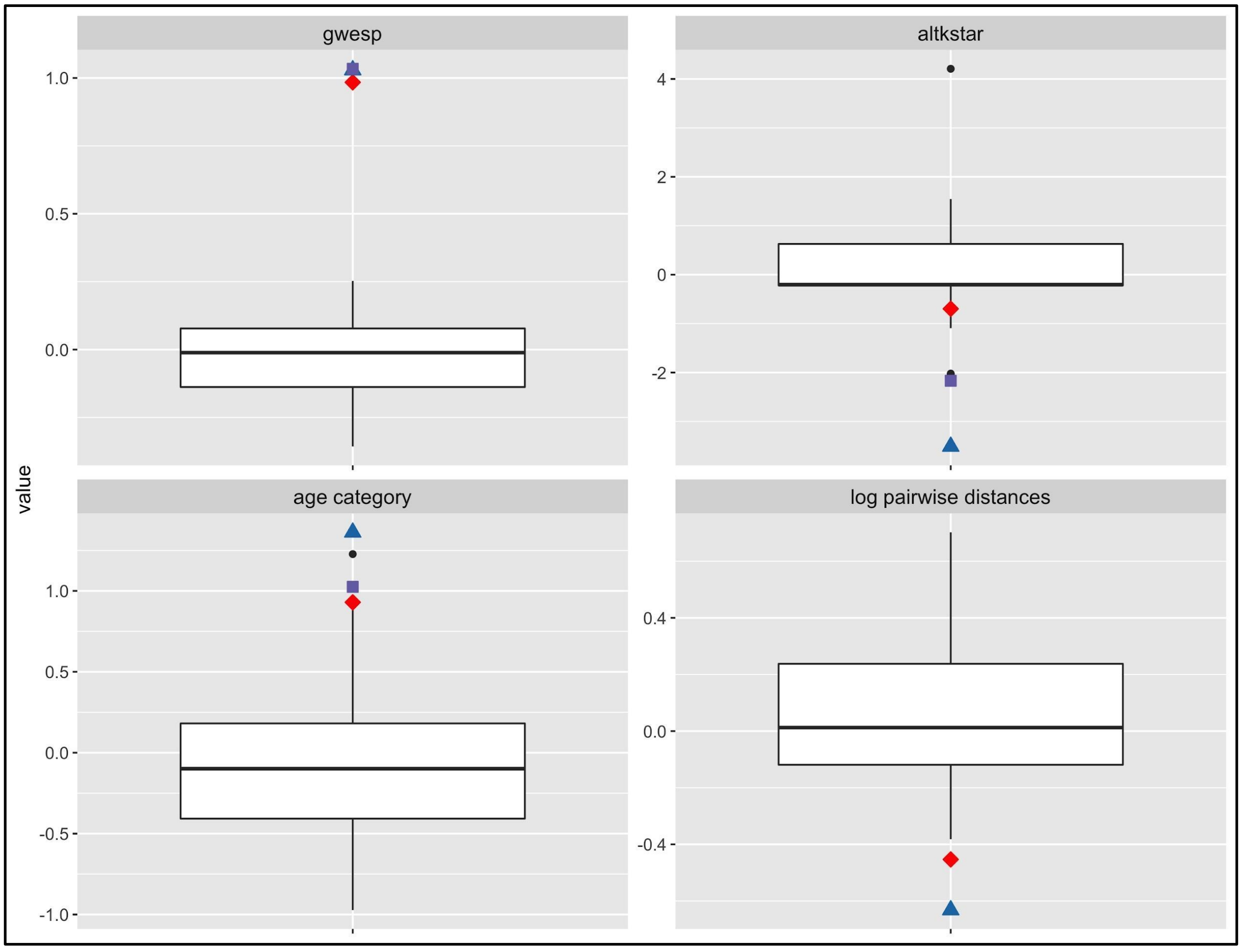

***Figure S4:*** *Fitting the predictors from the best ERGMs to random networks (based on the main FIV network) shows that all three best models give largely consistent coefficient estimates. Boxplots show coefficient estimates from 50 random networks. Red diamonds are estimates*

*from the main FIV network ERGM; blue triangles are estimates from the summary network with window overlap; purple squares are estimates from the summary network without window overlap. Note that the primary inconsistency between models is that the summary network without overlap did not identify log pairwise distances as a significant predictor.*

*FeLV spatial analyses*

As reported in the main text, SaTScan analysis of observed FeLV status found weak evidence of spatial clustering (two clusters detected, but not statistically significant with $p$=0.165 and 0.997, respectively; Figure S5).

279

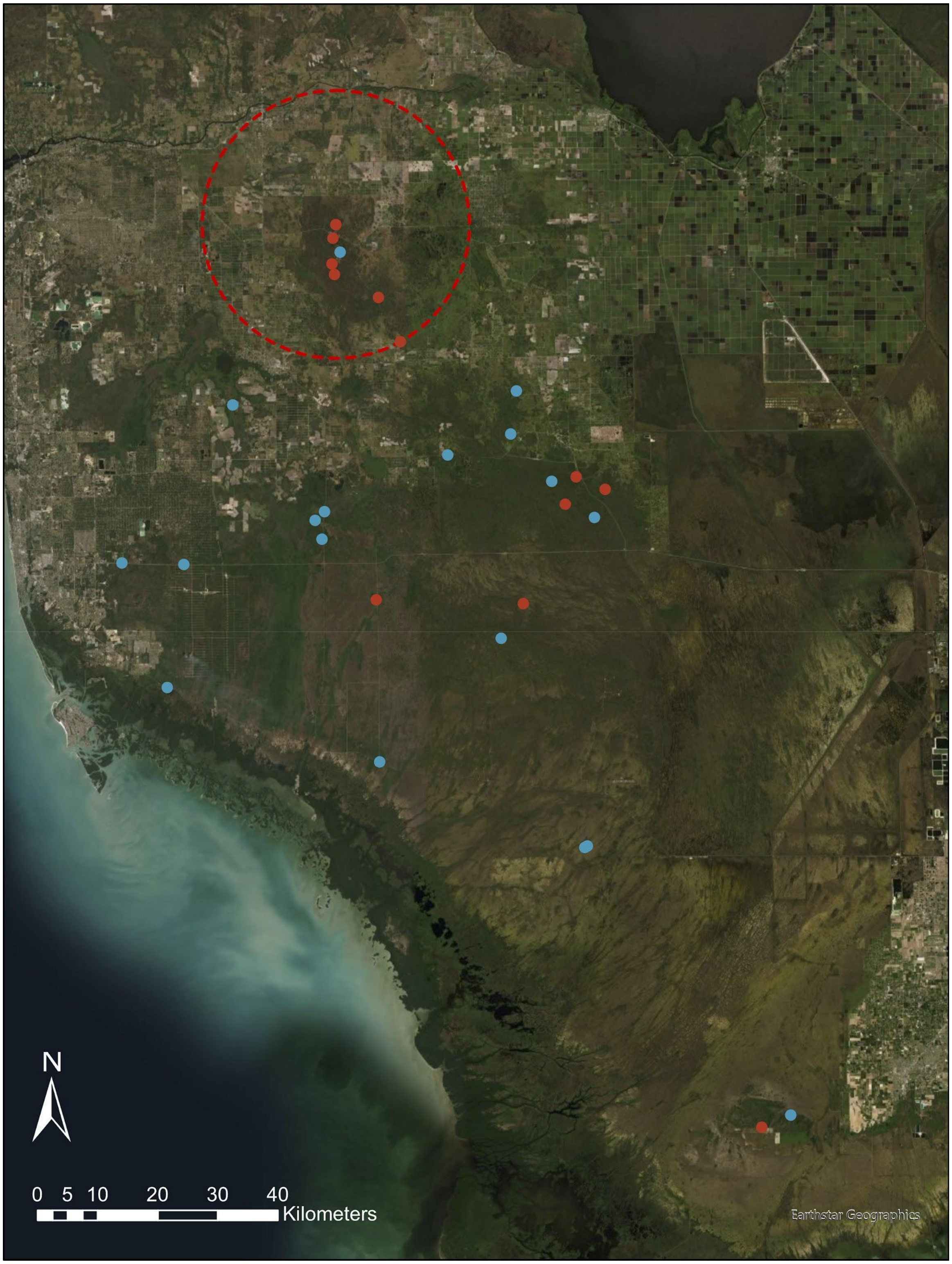

***Figure S5:*** *Map of observed panther centroid locations and FeLV status (red = qPCR positive, blue = qPCR negative). The red dashed circle shows the location and size of the top SaTScan cluster candidate, though this cluster was not considered statistically significant (p* = 0.165).

*FeLV simulations*

Main results of the generalized linear mixed model (GLMM) for FeLV predictive model performance (see main text) are given in Table 2 in the main text (homogeneous mixing model was reference group; for parameter set random intercepts: variance = 0.90; standard deviation = 0.95). While the FIV-based approach did not show statistically significant improvements in performance, it did trend toward best performance, having the highest number of “feasible” parameter sets (Figures S6, S7).

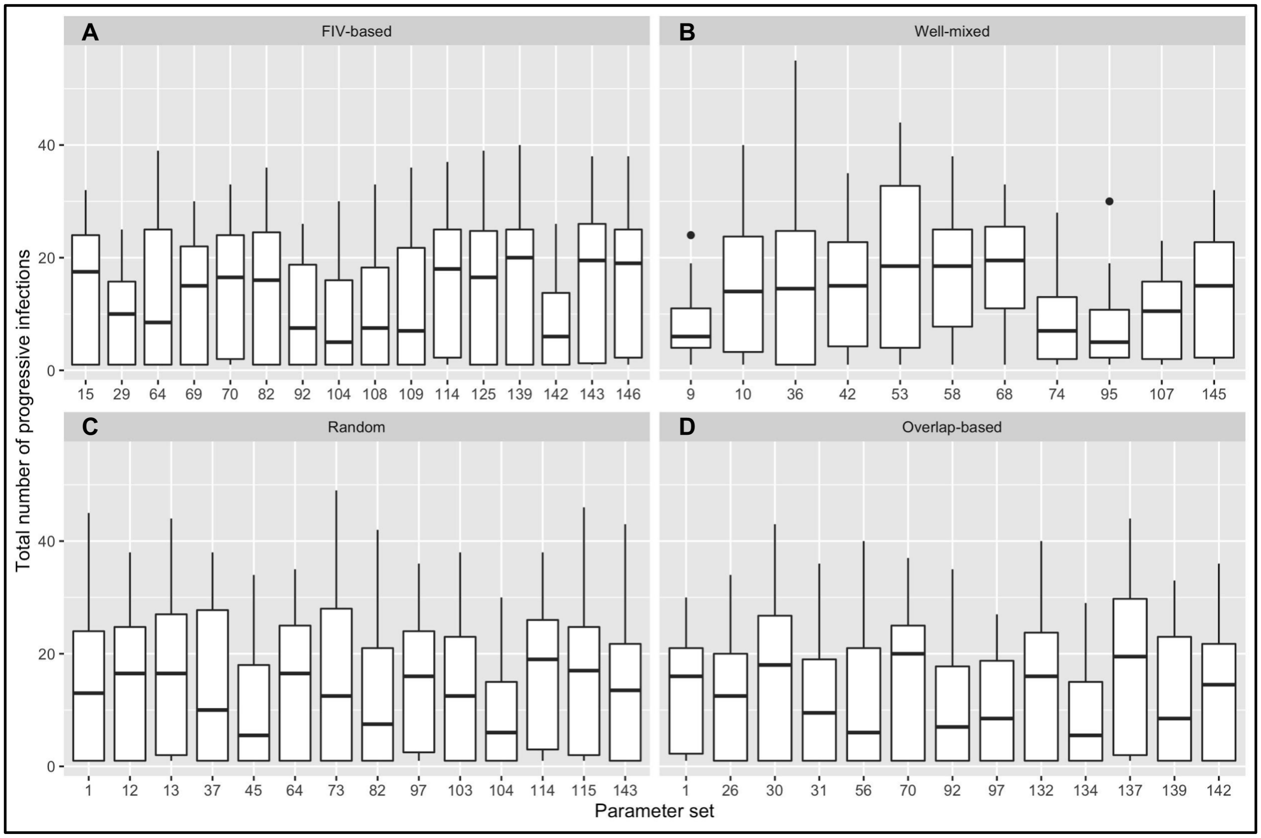

***Figure S6:*** *Boxplots of total number of progressive infections from parameter sets classified as "feasible." Results are shown for model types (A) FIV-based network, (B) well-mixed compartmental model, (C) random network, and (D) overlap-based network. Parameter set on the x-axis represents the unique parameter set drawn from our LHS sampling design; for example, set 1 for the FIV-based model type is identical to set 1 for random network model type, but feasible sets are not necessarily the same across model types.*

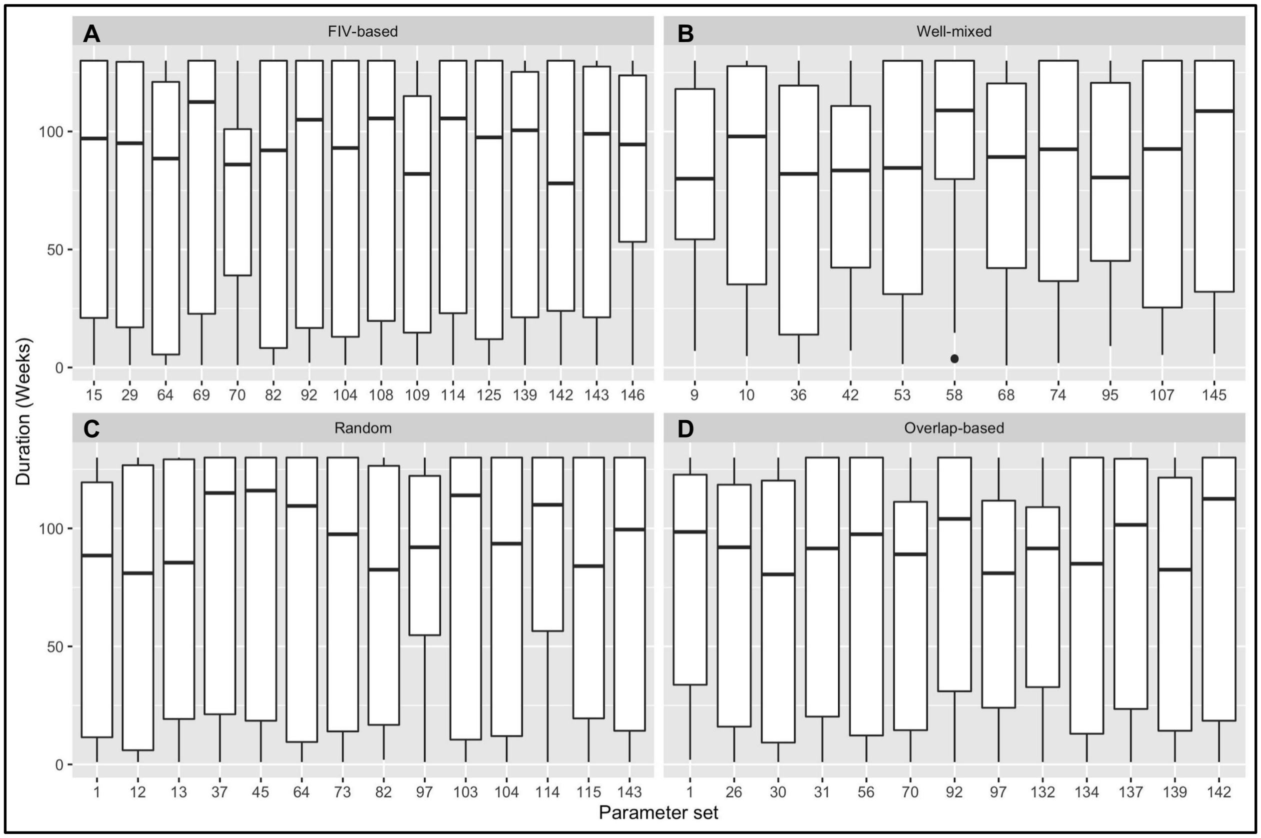

***Figure S7:*** *Boxplots of duration of simulated epidemics from parameter sets classified as "feasible." Results are shown for model types (A) FIV-based network, (B) well-mixed compartmental model, (C) random network, and (D) overlap-based network. Parameter set on the x-axis represents the unique parameter set drawn from our LHS sampling design; for example, set 1 for the random network model type is identical to set 1 for the overlap-based network model type, but feasible sets are not necessarily the same across model types.*

SaTScan results for simulated FeLV cases and controls are given in the main text. Cuzick and Edward's tests found evidence of global clustering of simulated FeLV cases with both the FIV and overlap-based models. However, for simulations with *p*-values less than or equal to 0.1, the FIV-based model was moderately more likely to capture the strength of global clustering (observed/expected test statistic, $T_k$, ratio) from the empirical FeLV data (Figure S8).

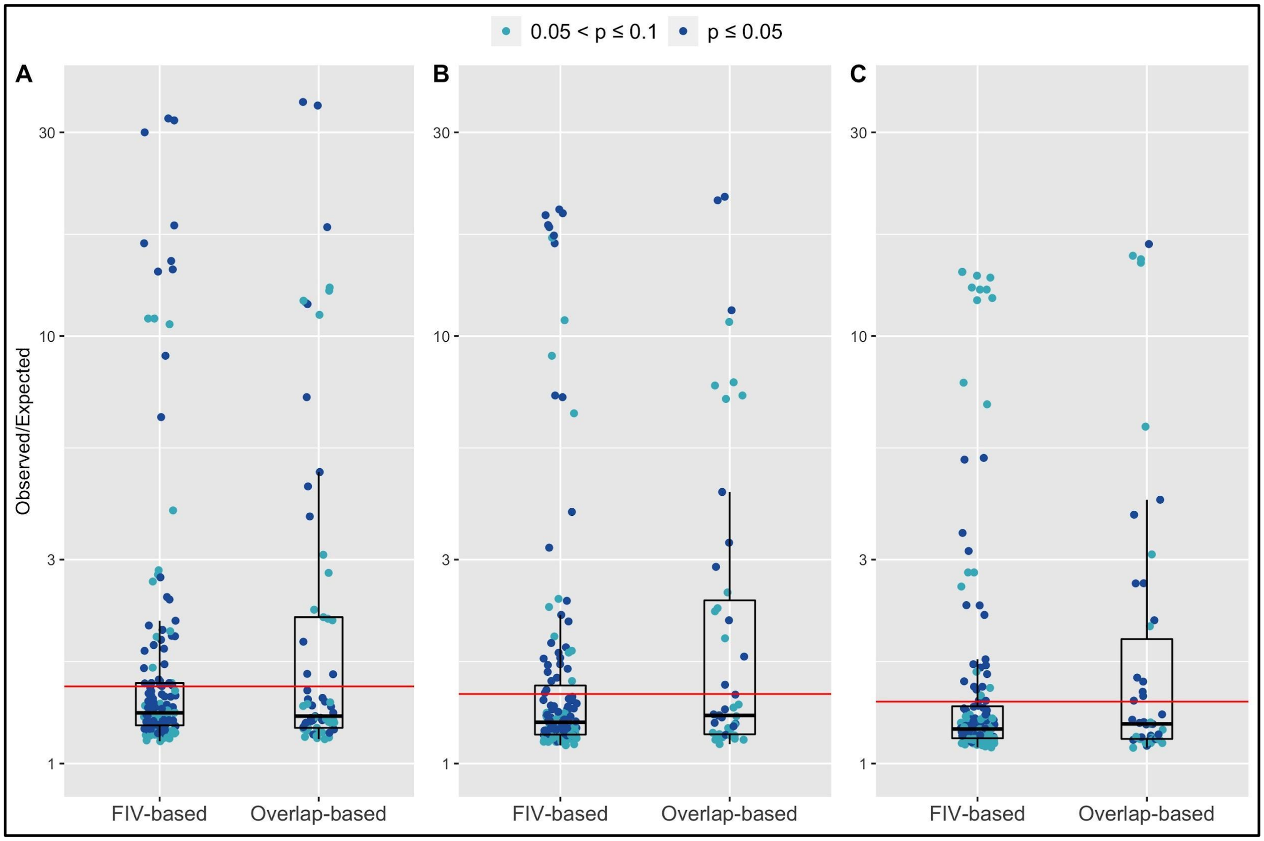

***Figure S8:*** *Observed/Expected test statistics ($T_k$) for Cuzick-Edwards tests performed with FIV and overlap-based predictions of FeLV transmission. Shown are results from feasible parameter set simulations in which Cuzick-Edwards test results had* p-*values less than or equal to 0.1. Plots represent the neighbor levels that demonstrated statistically significant clustering for the empirical FeLV data: (A) k = 3; (B) k = 5; (C) k = 7. The red horizontal line in all cases is the Observed/Expected $T_k$ ratio for the empirical FeLV data.*

In addition, to better understand the importance of FeLV transmission parameters in generating "feasible" results, we performed a *post hoc* random forest analysis for each model type, with "feasible" as a binary outcome for each parameter set (as in [22]). Predictors were the FeLV transmission transmission parameters, and data were split into 80% training/20% testing data sets. Because few parameter sets were categorized as feasible, we tested different resampling strategies for balancing the data. These included no resampling, down sampling, up sampling, up/down sampling, and SMOTE sampling (R package *DMwR*; [23]). The sampling protocol that produced the highest balanced accuracy was carried forward for analysis. In addition, we optimized hyper-parameters for the final random forest model. All random forests were performed using the *randomForest* package in R [24]. Across all model types, final random forest results tended to show poor balanced accuracy, low area under the curve (AUC; observed as low as AUC of 0.5) results, and were often inconsistent between repetitions (i.e. changes to training/testing data sets). Example random forest output for the FIV-based model is shown in Figures S9 and S10 for transparency, but should be interpreted with caution. Of particular note, however, was that C, the modifier shaping potential transmission from regressively infected individuals, had a strong tendency across model types to show best performance at C = 0.1 or 0.5 (Figure S11); this would support the possibility of low transmissibility of regressively infected individuals. See main text for further discussion of this finding.

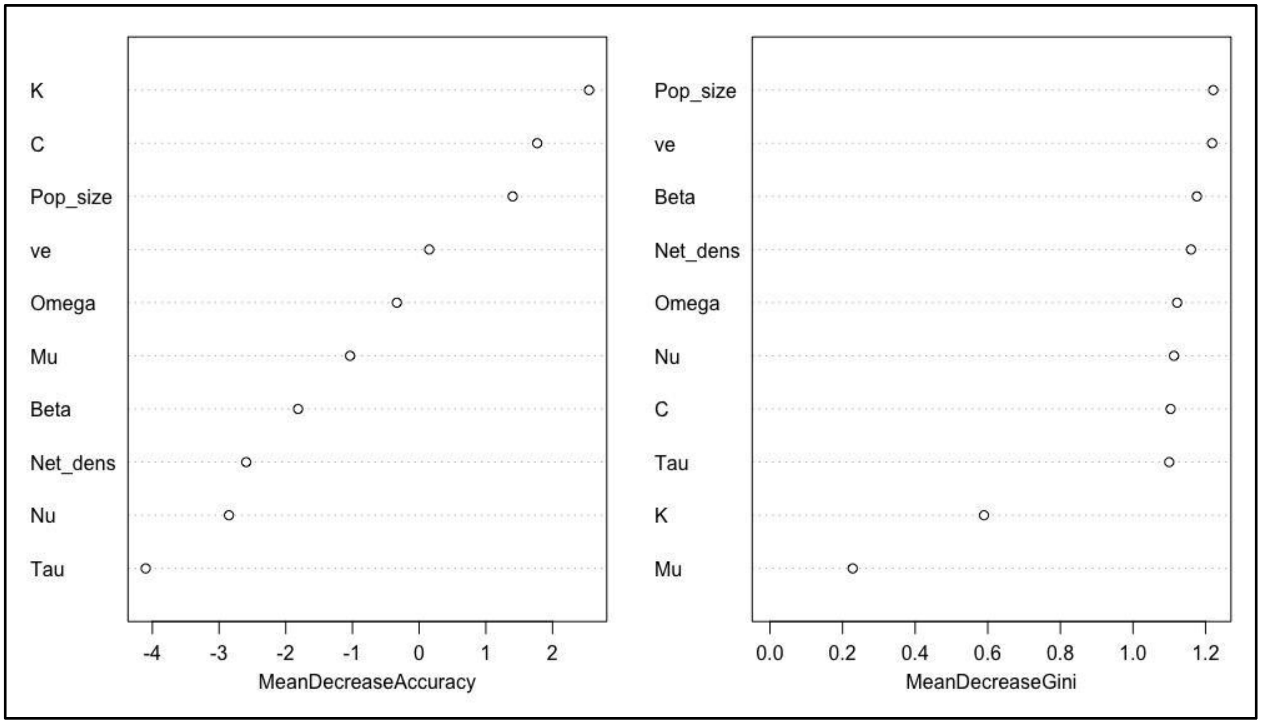

***Figure S9:*** *Variable importance plots for the FIV-based model. While AUC was 0.889 for this random forest analysis, results were inconsistent between random forests and should be interpreted with caution. Variable names are given on the x-axis (see Table S1). Mean decrease in accuracy scores is given on the x-axis in the left panel; mean decrease in Gini index on the x-axis in the right panel.*

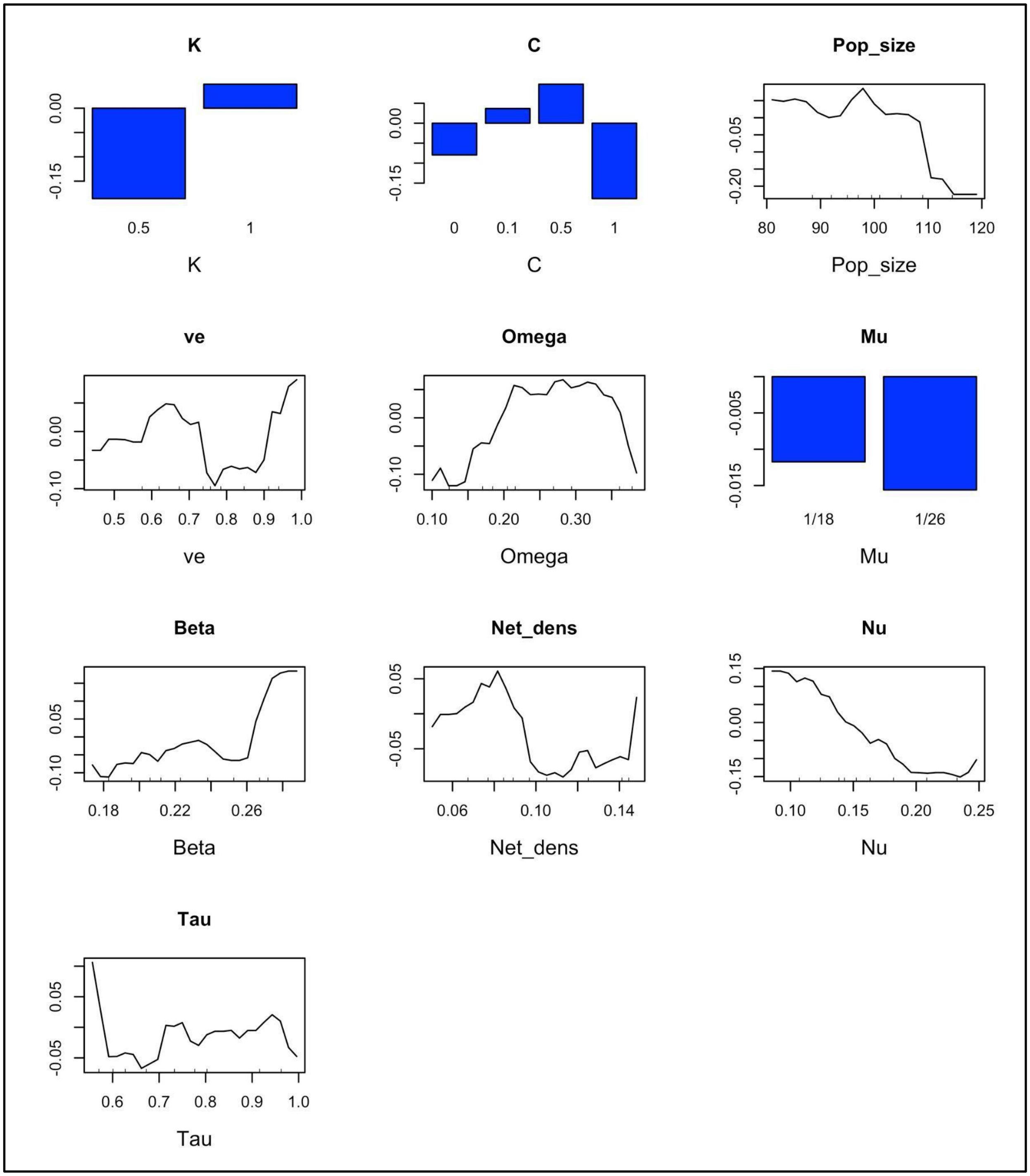

***Figure S10:*** *Partial dependence plots for the FIV-based model, ordered based on variable importance observable in Figure S9 (according to mean decrease in accuracy scores; highest importance in top left). While AUC was 0.889 for this random forest analysis, results were inconsistent between random forests and should be interpreted with caution. For example, note*

that partial dependence results differ quantitatively for the C parameter between this figure and Figure S11, though they are qualitatively consistent. Variable names are given in plot titles and x-axes (see Table S1).

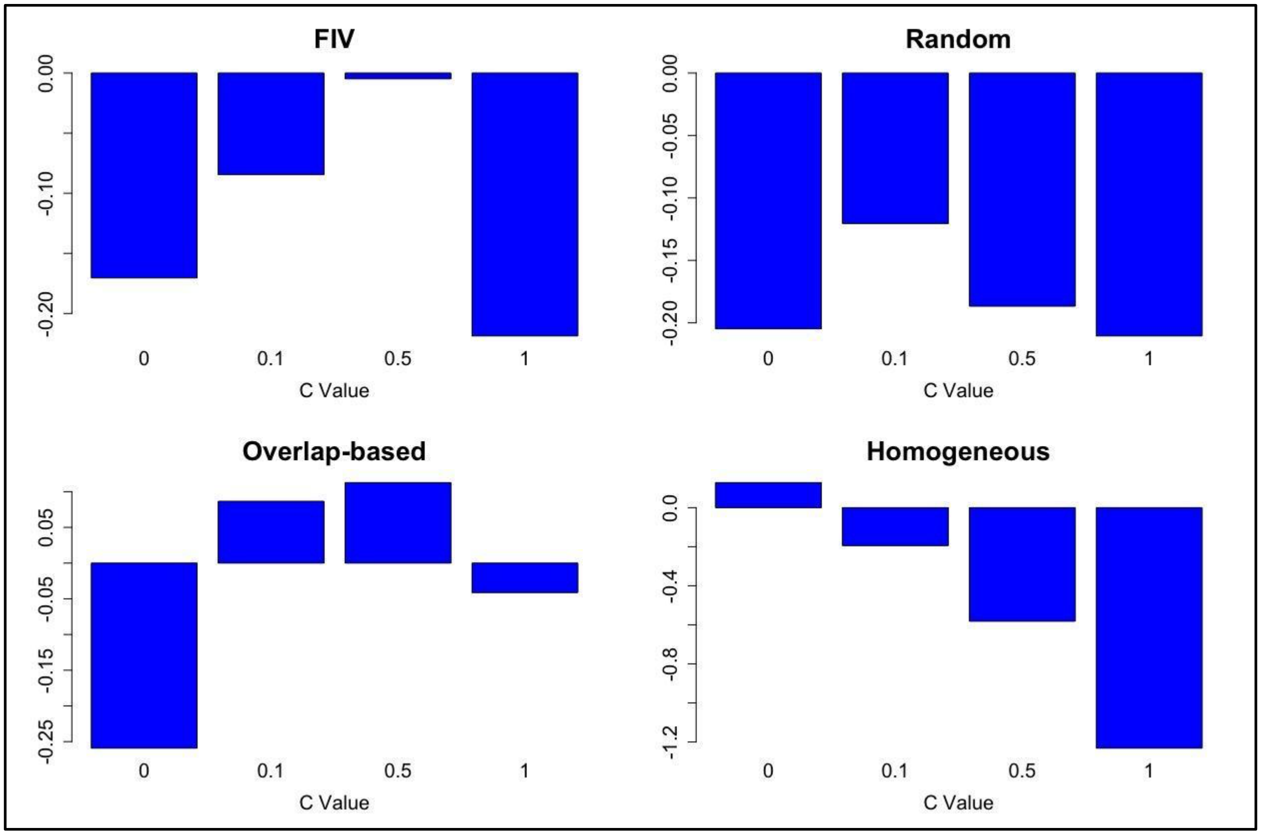

***Figure S11:*** *Partial dependence plots for the C parameter, which was a constant multiplier for probability of transmission from regressives, given effective contact. Because random forest analyses were sensitive to sampling, the plotted results are from random forest models in which the area under the curve was greater than or equal to 0.8. All models but homogeneous mixing showed at least some support for values of C greater than 0 but less than 1.*
